# Regression of solid breast tumours in mice by Newcastle disease virus is associated with production of apoptosis related-cytokines

**DOI:** 10.1101/374157

**Authors:** Umar Ahmad, Juraimi Raihan, Yoke Keong Yong, Zolkapli Eshak, Fauziah Othman, Aini Ideris

## Abstract

**Background:** Different strains of Newcastle disease virus (NDV) worldwide proved to have tumouricidal activity in several types of cancer cells. However, the possible anti-cancer activity of Malaysian NDV AF2240 strain and its mechanism of action remains unknown. The ability of cytokine-related apoptosis-inducing NDV AF2240 to treat breast cancer was investigated in the current study.

**Methods:** A total of 90 mice were used and divided into 15 groups, each group comprising of 6 mice. Tumour, body weight and mortality of the mice were determined throughout the experiment, to observe the effect of NDV and NDV+Tamoxifen treatments on the mice. In addition, the toxic effect of the treatments was determined through liver function test. In order to elucidate the involvement of cytokine production induced by NDV, a total of six cytokines, i.e. IL-6, IFN-γ, MCP-1, IL-10, IL12p70 and TNF-α were measured using cytometric bead array assay (plasma) and enzyme-linked immunosorbent spot (isolated splenocytes).

**Results:** The results demonstrated that 4T1 breast cancer cells in allotransplanted mice treated with AF2240 showed a noticeable inhibition in tumour growth and induce apoptotic-related cytokines.

**Conclusion:** NDV AF2240 suppression of breast tumour growth is associated with induction of apoptotic-related cytokines.

## 1. Background

To date, cancer remains the tenth leading causes of death worldwide. According to National Cancer Registry Report 2007, breast cancer is the top ranking malignancy that affects Malaysians followed by colorectal cancer[1]. There are a total of 3,292 new cases reported in the year 2007 and contributed 18.1% out of all the types of cancers in Malaysia[1]. However, the number of new cases are slightly less compared with the cases in 2003 (3738 cases reported) [2, 3] and the patient survival rate has also improved. This could be as a result of sensitization campaign, which increases awareness and educates citizen. In addition, early detection, development of advanced medical instruments and surgical techniques, and discovery of various new and effective anti-cancer agents may have resulted in the reduction in the number of cases.

There are various types of treatments available for breast cancer patients, including surgery, radiotherapy and chemotherapy. Also, numerous anti-cancer drugs, such as tamoxifen [4], trastuzumab [5] and bevacizumab [6], have been in the market since their discoveries. However, these drugs show many side effects. For instance, anastrozole that is used for the treatment of breast cancer patients after surgery leads to severe memory impairment [7] and long term use of tamoxifen increases the risk of endometrial cancer [8]. Due to this, tremendous efforts have been made by scientists around the world to explore and develop targeted drug systems with least or no side effects that will specifically target cancer cells without affecting normal healthy cells.

Recently, the trend of manipulating a virus to serve as an anti-cancer agent has been increasing [9, 10]. Among all types of viruses, oncolytic virus specifically targets cancer cell without causing excessive damage to non-cancerous cells [11]. Numerous oncolytic viruses from different family exhibit different mechanism of tumour selectivity. For instance, herpes simplex virus mutant, G207 from *herpesviridae* family has been used for the treatment of malignant glioma [12]. Mumps virus from the family of *paramyxoviridae* has been previously used against ovarian cancer [13], and Sindbis virus from *togaviridae* family has been proposed as a therapy for cervical and ovarian cancers [14].

Apart from all these viruses, oncolytic Newcastle disease virus (NDV) also has potential as an anti-cancer agent because of its inability to induce immune escape mechanism in mammalian cells[15]. Thus, it may be suitable as an immunologic adjuvant in a human cancer vaccine [16]. NDV belongs to *paramyxoviridae* family, and causes inflammation of respiratory tract, brain and gastrointestinal tract to poultry [17]. However, it only causes mild flu-like symptoms and conjunctivitis in human [18]. The most common strain that is found in Malaysia is NDV AF2440 [19]. It has been reported that NDV AF2240 was capable of stimulating apoptosis in breast cancer cells and the apoptotic effects were correlated with the production of pro-inflammatory cytokines in the tumour cells [20]. These effects are the first steps of immunotherapeutic effects of many crucial cytokines that help in combating breast tumour cells [21]. Several strains of NDV such as 73-T, HUJ, PV701 (MK107), MTH68, and Ulster have been shown to exhibit similar oncolytic properties as that of NDV AF2240 strain. Furthermore, additional exploration of the three Malaysian oncolytic NDV strains, AF2240, F and V4, have also been studied on different types of cancer cell lines in both *in vivo* and *in vitro* screening [22-24]. Of all these strains, only AF2240 (velogenic) was found to be more effective and showed better cytotoxic effect on in vitro MCF-7 cells as compared to the V4-UPM (lentogenic) strain [25]. Thus, AF2240 strain has the most significant anti-cancer activity and had proven to be relatively effective in suppressing tumour growth through apoptosis induction[22, 26]. Given this, here, we aim to investigate the possible apoptotic mechanism of NDV AF2240 against 4T1 breast cancer cells through cytokines induction.

## 2. Results

### 2.1. Effect of NDV on body weight and tumour weight in allotransplanted mice

All groups of mice showed significant increment in body weight compared with beginning stage (Table 1). On the other hand, the tumour in all groups increased in size, weight and no signs of regression at the end of the experiment except for groups CNDV8, 16, 32, 64, CNDV8+T and CNDV16+T (Table 1). The six groups mentioned above did not show any tumour growth at all. Thus, the percentage of inhibition was up to 100%. In contrast, groups CNDV32+T and CNDV64+T did not show any signs of inhibition, and it seems there is an enhanced tumour growth. CT group which was treated Tamoxifen, a reference drug, showed slightly weak tumour suppression of about 16.55%.

**Table 1:** Effect of VVNDV AF2240 and Tamoxifen in mortality rate, body and tumour weight of mice. Results expressed in mean ± S.D. \**p* < 0.05 compared with beginning body weight; # *p*< 0.05 compared with cancer control (CC). CT, allotransplanted + tamoxifen; CNDV, allotransplanted + virus; CNDV+T, allotransplanted + virus + Tamoxifen; NG, negative.

| Group | Animal number | | | Body weight ± SD | | Tumour weight (g) | Inhibition (%) |
| --- | --- | --- | --- | --- | --- | --- | --- |
|  | Beginning | End | Mortality rate (%) | Beginning | End |  |  |
| CC | 6 | 6 | 0 | 16.66 ± 0.70 | 21.26 ± 1.24 * | 1.45 ± 0.43 | - |
| CT | 6 | 6 | 0 | 16.48 ± 0.09 | 20.88 ± 0.79 * | 1.21 ± 0.41 | 16.55 |
| **CNDV** |  |  |  |  |  |  |  |
| 8HA NDV | 6 | 6 | 0 | 16.77 ± 0.11 | 19.49 ± 0.30 * | NG # | 100 |
| 16HA NDV | 6 | 6 | 0 | 16.64 ± 0.32 | 19.30 ± 0.06 * | NG # | 100 |
| 32HA NDV | 6 | 6 | 0 | 16.86 ± 0.10 | 19.45 ± 0.10 * | NG # | 100 |
| 64HA NDV | 6 | 6 | 0 | 16.54 ± 0.31 | 19.66 ± 0.13 * | NG # | 100 |
| **CNDV + T** |  |  |  |  |  |  |  |
| 8HA NDV | 6 | 6 | 0 | 16.93 ± 0.41 | 19.33 ± 0.16 * | NG # | 100 |
| 16HA NDV | 6 | 6 | 0 | 16.86 ± 0.14 | 19.50 ± 0.30 * | NG # | 100 |
| 32HA NDV | 6 | 6 | 0 | 16.87 ± 0.06 | 21.00 ± 0.40 * | 2.27 ± 0.28 # | -56.55 |
| 64HA NDV | 6 | 6 | 0 | 16.66 ± 0.06 | 20.59 ± 0.36 * | 3.64 ± 0.23 # | -151.03 |

### 2.2. Liver function test

The activity of serum AST, ALT and the level of total bilirubin for all the treated groups are shown in Table 2. The level of total bilirubin for all groups, except for CNDV32+T and CNDV 64+T groups, showed no significant difference compared with the normal control group. Both treatments indicated a markedly elevated level of total bilirubin. In the case of enzyme activities, cancer control (CC), reference drug (CT), CNDV16 and CNDV64+T significantly increased AST activity. This is especially apparent in the group treated with Tamoxifen (CT), which showed an increase one-fold higher than the normal control. On the other hand, groups CT, CNDV8+T, CNDV32+T and CNDV64+T showed a significant increment of ALT activity as compared to normal control.

**Table 2.** Activities of ALT, AST and the level of total bilirubin in different groups of mice.

| Liver Function Tests |  |  |  |
| --- | --- | --- | --- |
| Groups | Total Bilirubin [mg/dl] | Aspartate Transaminase (AST) [U/l] | Alanine Transaminase (ALT) [U/l] |
| NC | 3.43 ± 0.40 | 120.67 ± 1.53 | 36.00 ± 2.00 |
| CC | 2.13 ± 0.95 | 214.67 ± 1.53 * | 53.67 ± 1.53 |
| CT | 5.27 ± 0.59 | 265.00 ± 2.00 * | 95.67 ± 1.53 * |
| NDV8 | 4.43 ± 0.40 | 153.33 ± 1.53 | 38.67 ± 1.53 |
| NDV16 | 5.75 ± 0.35 | 125.33 ± 93.82 | 41.67 ± 1.53 |
| NDV32 | 4.87 ± 0.35 | 105.00 ± 1.00 | 47.00 ± 1.00 |
| NDV64 | 5.13 ± 0.67 | 115.00 ± 11.27 | 52.67 ± 2.51 |
| CNDV8 | 4.47 ± 0.35 | 98.67 ± 1.53 | 26.33 ± 1.15 |
| CNDV16 | 3.00 ± 0.50 | 173.33 ± 1.53 | 57.00 ± 1.00 |
| CNDV32 | 2.87 ± 0.35 | 121.00 ± 2.00 | 49.00 ± 1.00 |
| CNDV64 | 3.40 ± 0.40 | 153.67 ± 1.53 | 49.33 ± 0.58 |
| CNDV8+T | 2.77 ± 0.45 | 166.67 ± 56.62 | 103.00 ± 2.00 * |
| CNDV16+T | 2.43 ± 0.40 | 108.00 ± 1.00 | 56.33 ± 1.53 |
| CNDV32+T | 10.67 ± 4.72 * | 122.33 ± 2.52 | 160.33 ± 25.66 * |
| CNDV64+T | 14.33 ± 4.04 * | 170.67 ± 1.53 | 146.33 ± 32.52 * |

NC, normal control; CC, cancer control; CT, allotransplanted + Tamoxifen; NNDV, virus alone; CNDV, allotransplanted + virus; CNDV + T, allotransplanted + virus + Tamoxifen; n = 6. Data are shown as mean ± S.D. \* $p < 0.05$ compared with NC.

### 2.3. Cytokine analysis of the plasma/ serum by CBA assay

Plasma/serum from mice was collected at week 1, 2, 3 and 4 from each group setting and cytokine concentrations were measured by CBA assay. A total of six different cytokines were determined, i.e. IL-6, IFN-γ, MCP-1, IL-10, IL-12p70 and TNF-α. Among all the cytokines that were measured, the concentration of IL-6, MCP-1 and IL-12p70 in the group of cancer control (CC) were found to have a significant increase throughout the four weeks compared with normal control. In addition, CC group was found to produce the highest level of IL-6, MCP-1 and IL-12p70 (54.2 ± 8.7 pg/ml, 40.1 ± 2.4 pg/ml, 11.2 ± 2.9 pg/ml of, respectively), compared to normal control which are only 9.5 ± 1.2 pg/ml, 4.1 ± 0.1 pg/ml and 0.8 ± 0.1 pg/ml, respectively [Table 3(a), Table 4(a) and Table 5(a)]. On the other hand, CC, CT, CNDV8, 16, 32 and 64 significantly produced IL-10 throughout the four weeks. However, production of IL-10 by group CNDV 16 ceased in week 3 and 4 [Table 6(a)]. There is no expression of IL-10 in the group NDV8 – 64. Apart from that, CC, CT, CNDV16+T, CNDV32+T and CNDV64+T showed relatively high secretion of TNF-α throughout the experiment [Table 7(a)]. In the case of IFN-γ, all groups showed varying results in different week and consider relatively low production [Table 8(a)].

**Table 3:**
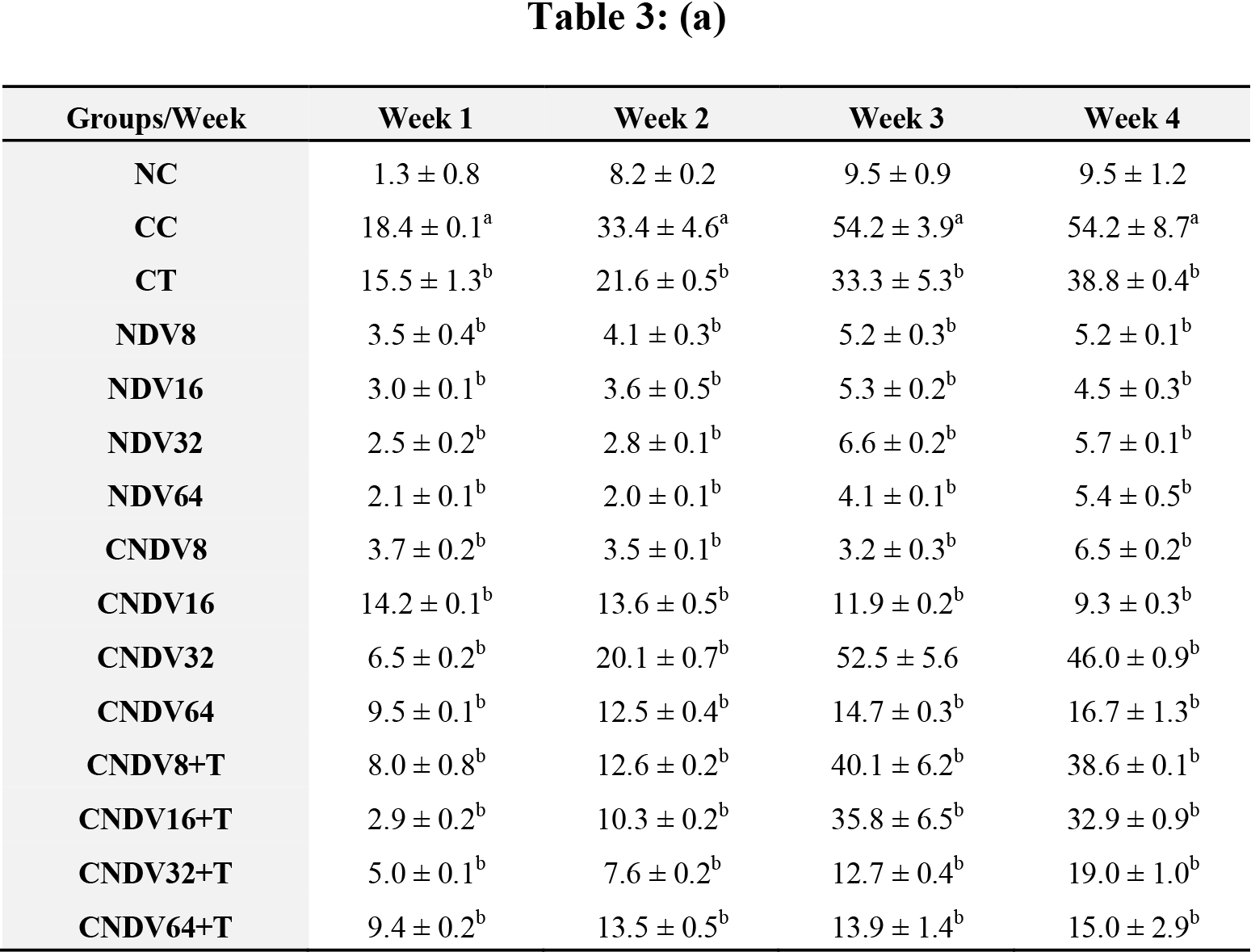
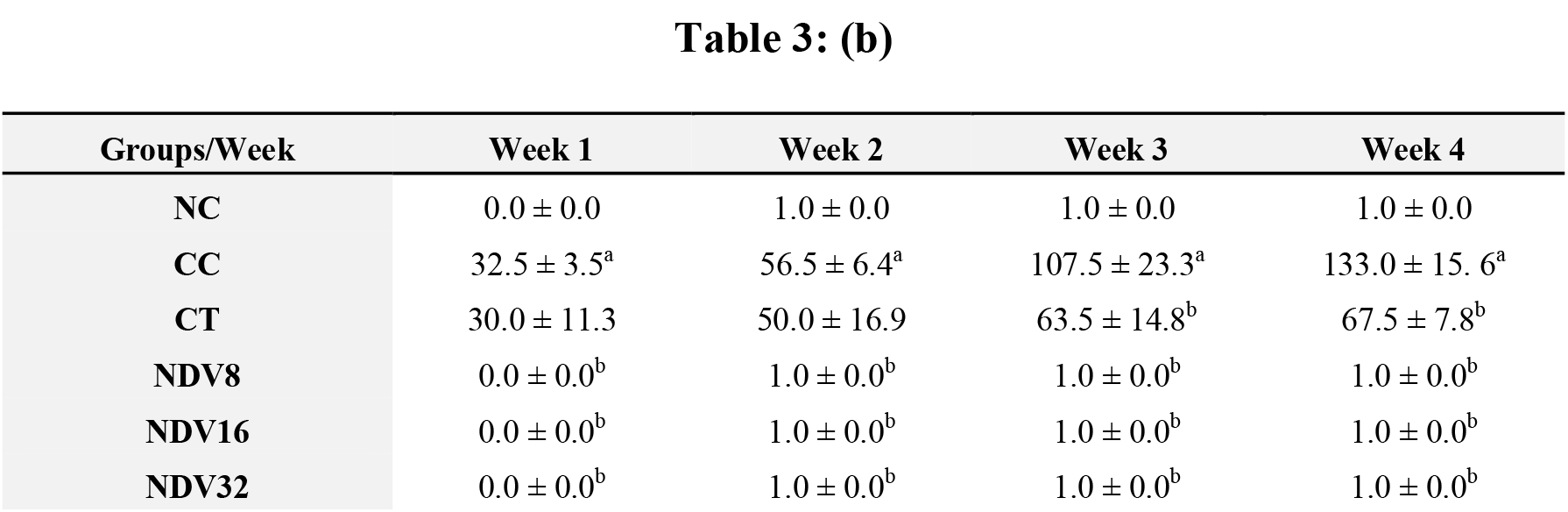
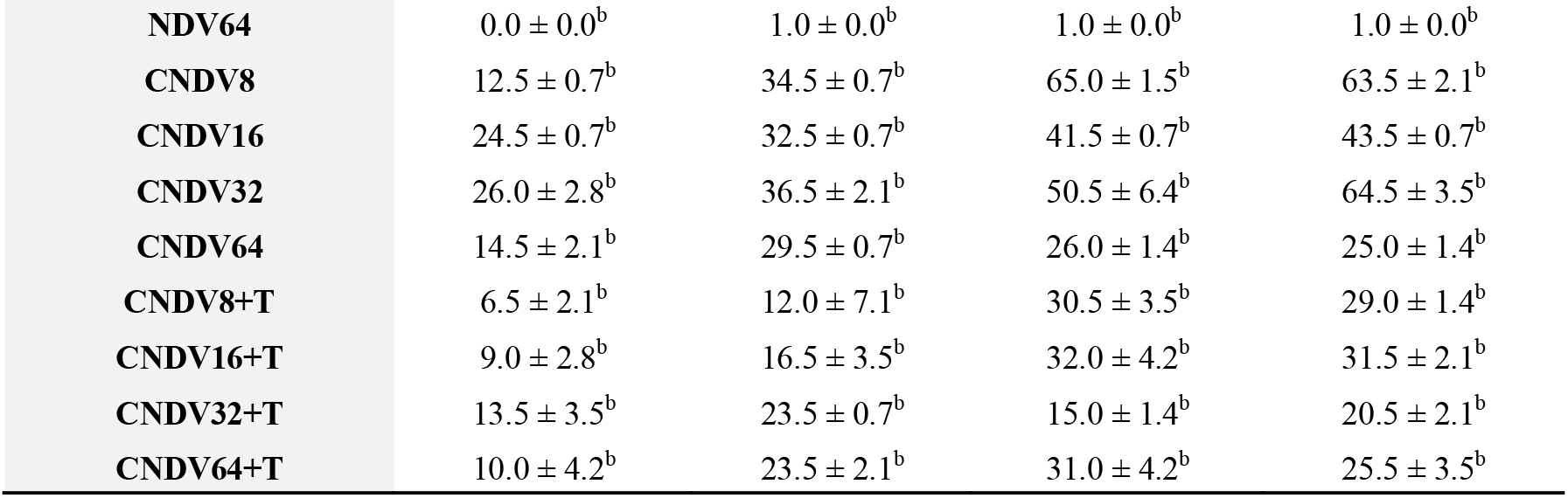
**(a)** Concentration (pg/ml) and **(b)** number spots of IL-6 in different groups of mice throughout four weeks experiment. NC: Normal Control; CC: Cancer Control; CT: Cancer + Tamoxifen; NDV: NDV virus alone; CNDV: Cancer and treated with NDV virus; CNDV + T: Cancer and treated with NDV and Tamoxifen. ^a^ *p* < 0.05 compared with NC; ^b^ *p* < 0.05 compared with CC.

| Groups/Week | Week 1 | Week 2 | Week 3 | Week 4 |
| --- | --- | --- | --- | --- |
| NC | 1.3 ± 0.8 | 8.2 ± 0.2 | 9.5 ± 0.9 | 9.5 ± 1.2 |
| CC | 18.4 ± 0.1<sup>a</sup> | 33.4 ± 4.6<sup>a</sup> | 54.2 ± 3.9<sup>a</sup> | 54.2 ± 8.7<sup>a</sup> |
| CT | 15.5 ± 1.3<sup>b</sup> | 21.6 ± 0.5<sup>b</sup> | 33.3 ± 5.3<sup>b</sup> | 38.8 ± 0.4<sup>b</sup> |
| NDV8 | 3.5 ± 0.4<sup>b</sup> | 4.1 ± 0.3<sup>b</sup> | 5.2 ± 0.3<sup>b</sup> | 5.2 ± 0.1<sup>b</sup> |
| NDV16 | 3.0 ± 0.1<sup>b</sup> | 3.6 ± 0.5<sup>b</sup> | 5.3 ± 0.2<sup>b</sup> | 4.5 ± 0.3<sup>b</sup> |
| NDV32 | 2.5 ± 0.2<sup>b</sup> | 2.8 ± 0.1<sup>b</sup> | 6.6 ± 0.2<sup>b</sup> | 5.7 ± 0.1<sup>b</sup> |
| NDV64 | 2.1 ± 0.1<sup>b</sup> | 2.0 ± 0.1<sup>b</sup> | 4.1 ± 0.1<sup>b</sup> | 5.4 ± 0.5<sup>b</sup> |
| CNDV8 | 3.7 ± 0.2<sup>b</sup> | 3.5 ± 0.1<sup>b</sup> | 3.2 ± 0.3<sup>b</sup> | 6.5 ± 0.2<sup>b</sup> |
| CNDV16 | 14.2 ± 0.1<sup>b</sup> | 13.6 ± 0.5<sup>b</sup> | 11.9 ± 0.2<sup>b</sup> | 9.3 ± 0.3<sup>b</sup> |
| CNDV32 | 6.5 ± 0.2<sup>b</sup> | 20.1 ± 0.7<sup>b</sup> | 52.5 ± 5.6 | 46.0 ± 0.9<sup>b</sup> |
| CNDV64 | 9.5 ± 0.1<sup>b</sup> | 12.5 ± 0.4<sup>b</sup> | 14.7 ± 0.3<sup>b</sup> | 16.7 ± 1.3<sup>b</sup> |
| CNDV8+T | 8.0 ± 0.8<sup>b</sup> | 12.6 ± 0.2<sup>b</sup> | 40.1 ± 6.2<sup>b</sup> | 38.6 ± 0.1<sup>b</sup> |
| CNDV16+T | 2.9 ± 0.2<sup>b</sup> | 10.3 ± 0.2<sup>b</sup> | 35.8 ± 6.5<sup>b</sup> | 32.9 ± 0.9<sup>b</sup> |
| CNDV32+T | 5.0 ± 0.1<sup>b</sup> | 7.6 ± 0.2<sup>b</sup> | 12.7 ± 0.4<sup>b</sup> | 19.0 ± 1.0<sup>b</sup> |
| CNDV64+T | 9.4 ± 0.2<sup>b</sup> | 13.5 ± 0.5<sup>b</sup> | 13.9 ± 1.4<sup>b</sup> | 15.0 ± 2.9<sup>b</sup> |
| NC | 0.0 ± 0.0 | 1.0 ± 0.0 | 1.0 ± 0.0 | 1.0 ± 0.0 |
| CC | 32.5 ± 3.5<sup>a</sup> | 56.5 ± 6.4<sup>a</sup> | 107.5 ± 23.3<sup>a</sup> | 133.0 ± 15. 6<sup>a</sup> |
| CT | 30.0 ± 11.3 | 50.0 ± 16.9 | 63.5 ± 14.8<sup>b</sup> | 67.5 ± 7.8<sup>b</sup> |
| NDV8 | 0.0 ± 0.0<sup>b</sup> | 1.0 ± 0.0<sup>b</sup> | 1.0 ± 0.0<sup>b</sup> | 1.0 ± 0.0<sup>b</sup> |
| NDV16 | 0.0 ± 0.0<sup>b</sup> | 1.0 ± 0.0<sup>b</sup> | 1.0 ± 0.0<sup>b</sup> | 1.0 ± 0.0<sup>b</sup> |
| NDV32 | 0.0 ± 0.0<sup>b</sup> | 1.0 ± 0.0<sup>b</sup> | 1.0 ± 0.0<sup>b</sup> | 1.0 ± 0.0<sup>b</sup> |
| **NDV64** | 0.0 ± 0.0<sup>b</sup> | 1.0 ± 0.0<sup>b</sup> | 1.0 ± 0.0<sup>b</sup> | 1.0 ± 0.0<sup>b</sup> |
| **CNDV8** | 12.5 ± 0.7<sup>b</sup> | 34.5 ± 0.7<sup>b</sup> | 65.0 ± 1.5<sup>b</sup> | 63.5 ± 2.1<sup>b</sup> |
| **CNDV16** | 24.5 ± 0.7<sup>b</sup> | 32.5 ± 0.7<sup>b</sup> | 41.5 ± 0.7<sup>b</sup> | 43.5 ± 0.7<sup>b</sup> |
| **CNDV32** | 26.0 ± 2.8<sup>b</sup> | 36.5 ± 2.1<sup>b</sup> | 50.5 ± 6.4<sup>b</sup> | 64.5 ± 3.5<sup>b</sup> |
| **CNDV64** | 14.5 ± 2.1<sup>b</sup> | 29.5 ± 0.7<sup>b</sup> | 26.0 ± 1.4<sup>b</sup> | 25.0 ± 1.4<sup>b</sup> |
| **CNDV8+T** | 6.5 ± 2.1<sup>b</sup> | 12.0 ± 7.1<sup>b</sup> | 30.5 ± 3.5<sup>b</sup> | 29.0 ± 1.4<sup>b</sup> |
| **CNDV16+T** | 9.0 ± 2.8<sup>b</sup> | 16.5 ± 3.5<sup>b</sup> | 32.0 ± 4.2<sup>b</sup> | 31.5 ± 2.1<sup>b</sup> |
| **CNDV32+T** | 13.5 ± 3.5<sup>b</sup> | 23.5 ± 0.7<sup>b</sup> | 15.0 ± 1.4<sup>b</sup> | 20.5 ± 2.1<sup>b</sup> |
| **CNDV64+T** | 10.0 ± 4.2<sup>b</sup> | 23.5 ± 2.1<sup>b</sup> | 31.0 ± 4.2<sup>b</sup> | 25.5 ± 3.5<sup>b</sup> |

**Table 4:**
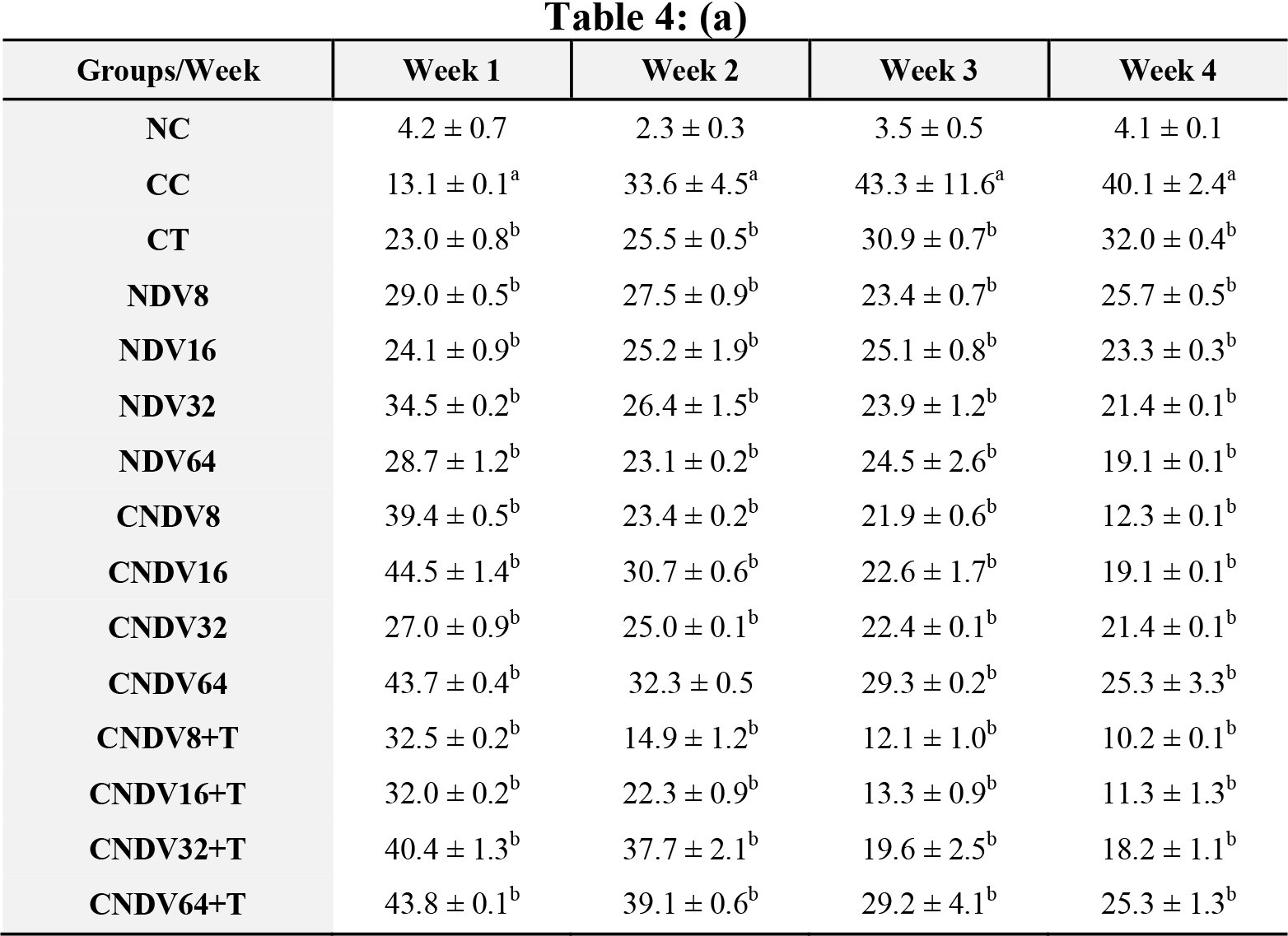
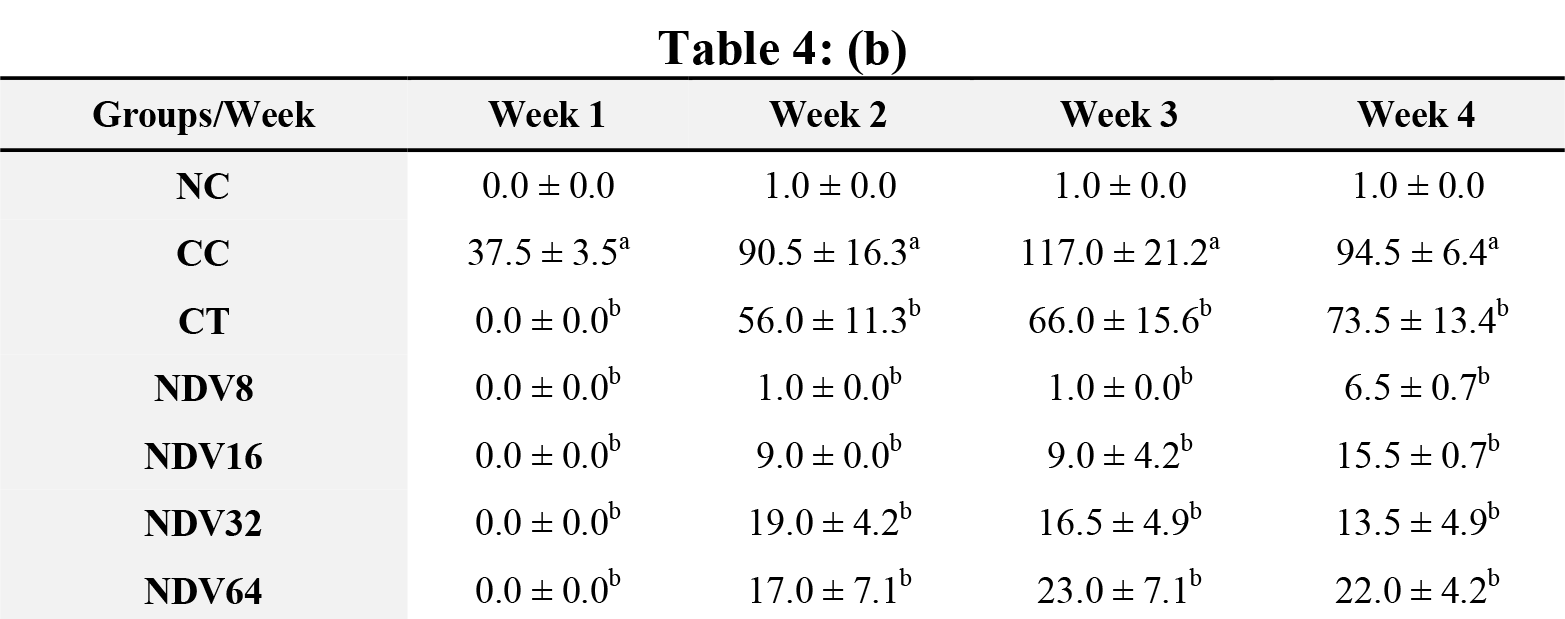
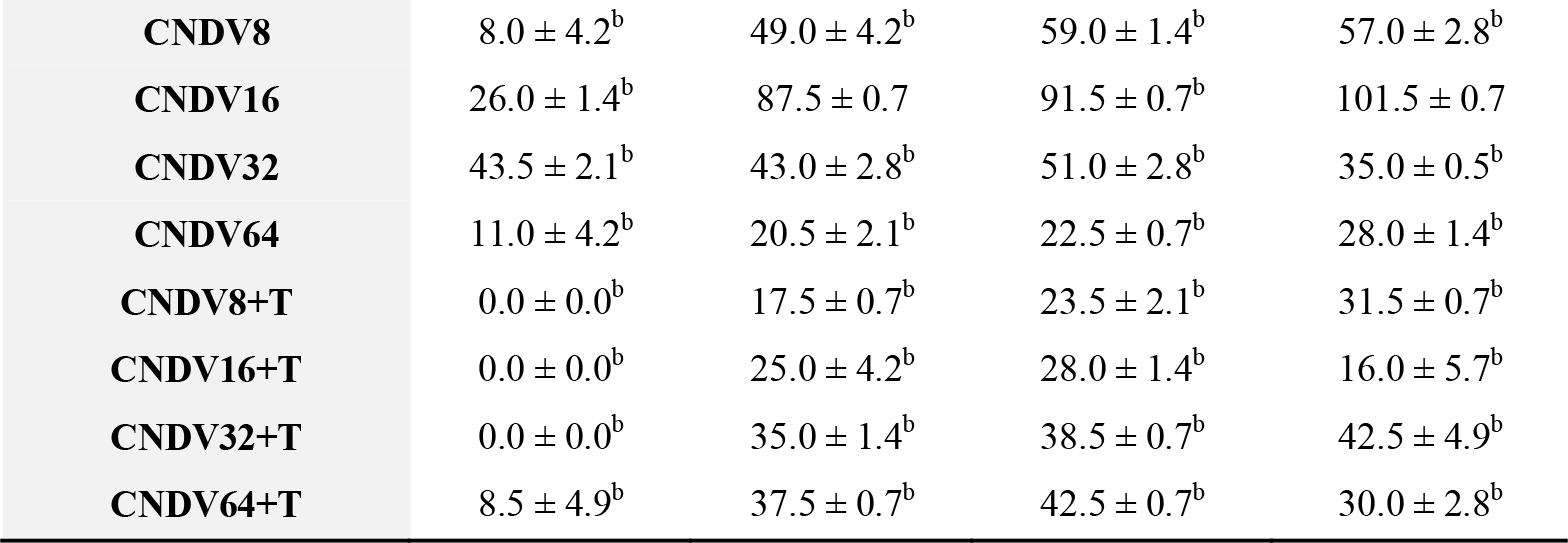
**(a)** Concentration (pg/ml) and **(b)** number spots of MCP-1 in different groups of mice throughout four weeks experiment. NC: Normal Control; CC: Cancer Control; CT: Cancer + Tamoxifen; NDV: NDV virus alone; CNDV: Cancer and treated with NDV virus; CNDV + T: Cancer and treated with NDV and Tamoxifen. ^a^ *p* < 0.05 compared with NC; ^b^ *p* < 0.05 compared with CC.

| Groups/Week | Week 1 | Week 2 | Week 3 | Week 4 |
| --- | --- | --- | --- | --- |
| **NC** | 4.2 ± 0.7 | 2.3 ± 0.3 | 3.5 ± 0.5 | 4.1 ± 0.1 |
| **CC** | 13.1 ± 0.1<sup>a</sup> | 33.6 ± 4.5<sup>a</sup> | 43.3 ± 11.6<sup>a</sup> | 40.1 ± 2.4<sup>a</sup> |
| **CT** | 23.0 ± 0.8<sup>b</sup> | 25.5 ± 0.5<sup>b</sup> | 30.9 ± 0.7<sup>b</sup> | 32.0 ± 0.4<sup>b</sup> |
| **NDV8** | 29.0 ± 0.5<sup>b</sup> | 27.5 ± 0.9<sup>b</sup> | 23.4 ± 0.7<sup>b</sup> | 25.7 ± 0.5<sup>b</sup> |
| **NDV16** | 24.1 ± 0.9<sup>b</sup> | 25.2 ± 1.9<sup>b</sup> | 25.1 ± 0.8<sup>b</sup> | 23.3 ± 0.3<sup>b</sup> |
| **NDV32** | 34.5 ± 0.2<sup>b</sup> | 26.4 ± 1.5<sup>b</sup> | 23.9 ± 1.2<sup>b</sup> | 21.4 ± 0.1<sup>b</sup> |
| **NDV64** | 28.7 ± 1.2<sup>b</sup> | 23.1 ± 0.2<sup>b</sup> | 24.5 ± 2.6<sup>b</sup> | 19.1 ± 0.1<sup>b</sup> |
| **CNDV8** | 39.4 ± 0.5<sup>b</sup> | 23.4 ± 0.2<sup>b</sup> | 21.9 ± 0.6<sup>b</sup> | 12.3 ± 0.1<sup>b</sup> |
| **CNDV16** | 44.5 ± 1.4<sup>b</sup> | 30.7 ± 0.6<sup>b</sup> | 22.6 ± 1.7<sup>b</sup> | 19.1 ± 0.1<sup>b</sup> |
| **CNDV32** | 27.0 ± 0.9<sup>b</sup> | 25.0 ± 0.1<sup>b</sup> | 22.4 ± 0.1<sup>b</sup> | 21.4 ± 0.1<sup>b</sup> |
| **CNDV64** | 43.7 ± 0.4<sup>b</sup> | 32.3 ± 0.5 | 29.3 ± 0.2<sup>b</sup> | 25.3 ± 3.3<sup>b</sup> |
| **CNDV8+T** | 32.5 ± 0.2<sup>b</sup> | 14.9 ± 1.2<sup>b</sup> | 12.1 ± 1.0<sup>b</sup> | 10.2 ± 0.1<sup>b</sup> |
| **CNDV16+T** | 32.0 ± 0.2<sup>b</sup> | 22.3 ± 0.9<sup>b</sup> | 13.3 ± 0.9<sup>b</sup> | 11.3 ± 1.3<sup>b</sup> |
| **CNDV32+T** | 40.4 ± 1.3<sup>b</sup> | 37.7 ± 2.1<sup>b</sup> | 19.6 ± 2.5<sup>b</sup> | 18.2 ± 1.1<sup>b</sup> |
| **CNDV64+T** | 43.8 ± 0.1<sup>b</sup> | 39.1 ± 0.6<sup>b</sup> | 29.2 ± 4.1<sup>b</sup> | 25.3 ± 1.3<sup>b</sup> |
| **NC** | 0.0 ± 0.0 | 1.0 ± 0.0 | 1.0 ± 0.0 | 1.0 ± 0.0 |
| **CC** | 37.5 ± 3.5<sup>a</sup> | 90.5 ± 16.3<sup>a</sup> | 117.0 ± 21.2<sup>a</sup> | 94.5 ± 6.4<sup>a</sup> |
| **CT** | 0.0 ± 0.0<sup>b</sup> | 56.0 ± 11.3<sup>b</sup> | 66.0 ± 15.6<sup>b</sup> | 73.5 ± 13.4<sup>b</sup> |
| **NDV8** | 0.0 ± 0.0<sup>b</sup> | 1.0 ± 0.0<sup>b</sup> | 1.0 ± 0.0<sup>b</sup> | 6.5 ± 0.7<sup>b</sup> |
| **NDV16** | 0.0 ± 0.0<sup>b</sup> | 9.0 ± 0.0<sup>b</sup> | 9.0 ± 4.2<sup>b</sup> | 15.5 ± 0.7<sup>b</sup> |
| **NDV32** | 0.0 ± 0.0<sup>b</sup> | 19.0 ± 4.2<sup>b</sup> | 16.5 ± 4.9<sup>b</sup> | 13.5 ± 4.9<sup>b</sup> |
| **NDV64** | 0.0 ± 0.0<sup>b</sup> | 17.0 ± 7.1<sup>b</sup> | 23.0 ± 7.1<sup>b</sup> | 22.0 ± 4.2<sup>b</sup> |
| **CNDV8** | 8.0 ± 4.2<sup>b</sup> | 49.0 ± 4.2<sup>b</sup> | 59.0 ± 1.4<sup>b</sup> | 57.0 ± 2.8<sup>b</sup> |
| **CNDV16** | 26.0 ± 1.4<sup>b</sup> | 87.5 ± 0.7 | 91.5 ± 0.7<sup>b</sup> | 101.5 ± 0.7 |
| **CNDV32** | 43.5 ± 2.1<sup>b</sup> | 43.0 ± 2.8<sup>b</sup> | 51.0 ± 2.8<sup>b</sup> | 35.0 ± 0.5<sup>b</sup> |
| **CNDV64** | 11.0 ± 4.2<sup>b</sup> | 20.5 ± 2.1<sup>b</sup> | 22.5 ± 0.7<sup>b</sup> | 28.0 ± 1.4<sup>b</sup> |
| **CNDV8+T** | 0.0 ± 0.0<sup>b</sup> | 17.5 ± 0.7<sup>b</sup> | 23.5 ± 2.1<sup>b</sup> | 31.5 ± 0.7<sup>b</sup> |
| **CNDV16+T** | 0.0 ± 0.0<sup>b</sup> | 25.0 ± 4.2<sup>b</sup> | 28.0 ± 1.4<sup>b</sup> | 16.0 ± 5.7<sup>b</sup> |
| **CNDV32+T** | 0.0 ± 0.0<sup>b</sup> | 35.0 ± 1.4<sup>b</sup> | 38.5 ± 0.7<sup>b</sup> | 42.5 ± 4.9<sup>b</sup> |
| **CNDV64+T** | 8.5 ± 4.9<sup>b</sup> | 37.5 ± 0.7<sup>b</sup> | 42.5 ± 0.7<sup>b</sup> | 30.0 ± 2.8<sup>b</sup> |

**Table 5:**
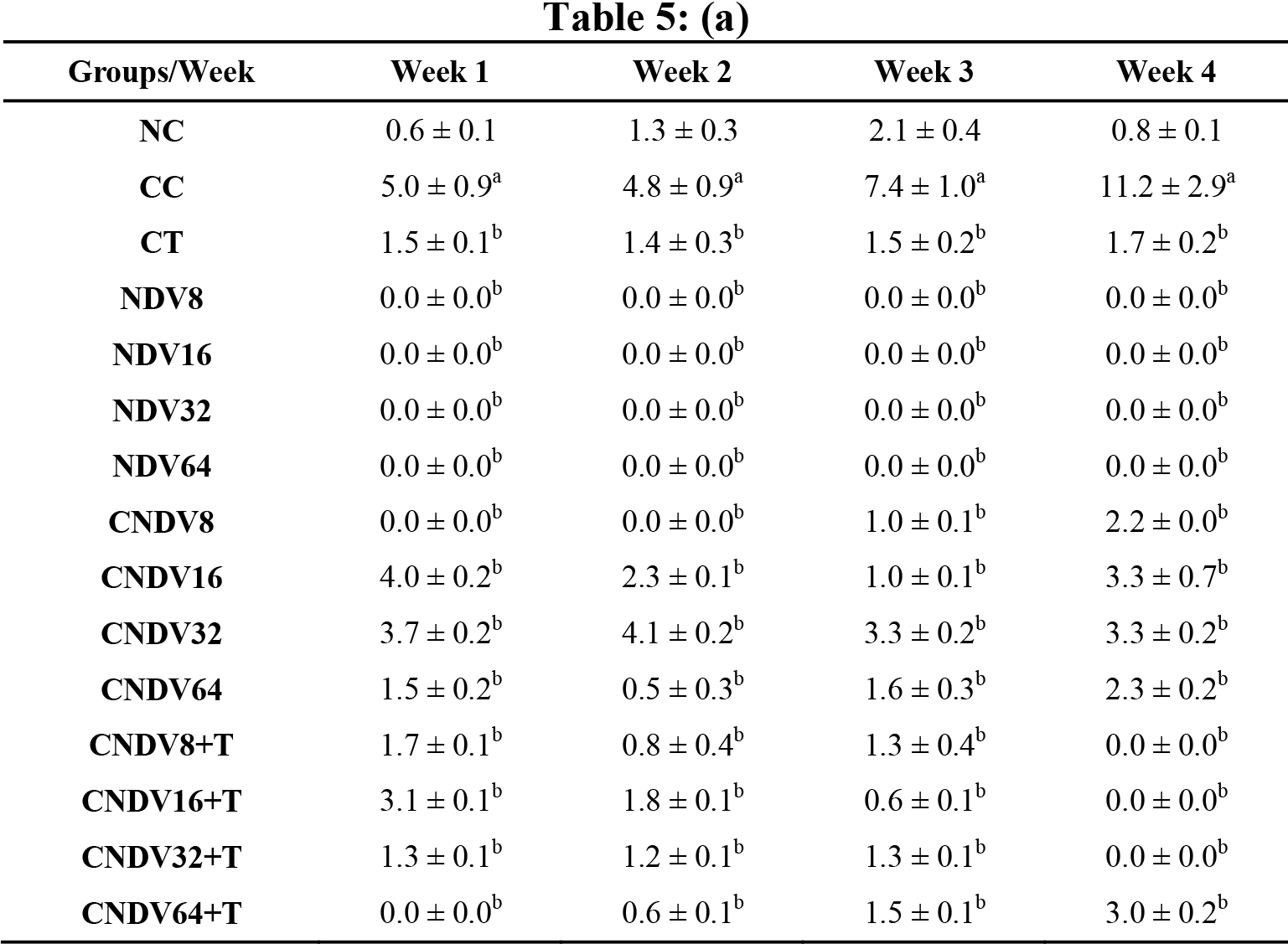
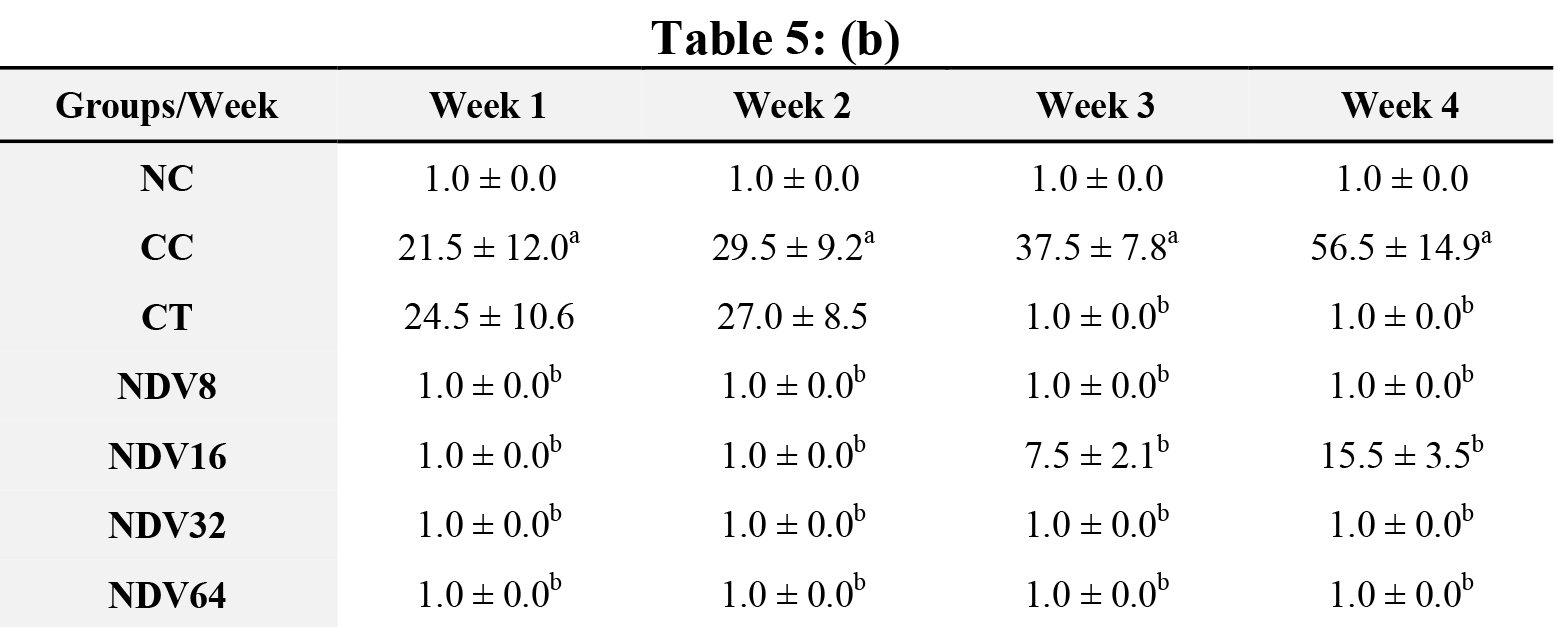
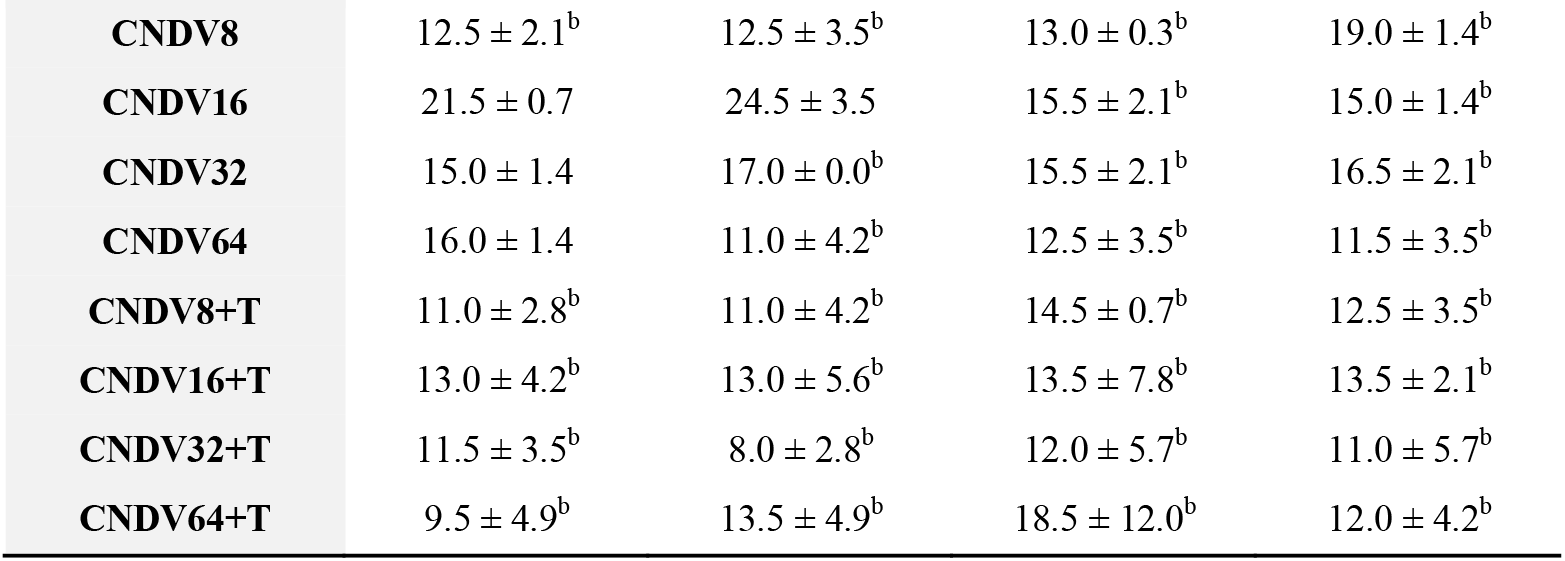
**(a)** Concentration (pg/ml) and **(b)** number spots of IL-12 in different groups of mice throughout four weeks experiment. NC: Normal Control; CC: Cancer Control; CT: Cancer + Tamoxifen; NDV: NDV virus alone; CNDV: Cancer and treated with NDV virus; CNDV + T: Cancer and treated with NDV and Tamoxifen. ^a^ *p* < 0.05 compared with NC; ^b^ *p* < 0.05 compared with CC.

| Groups/Week | Week 1 | Week 2 | Week 3 | Week 4 |
| --- | --- | --- | --- | --- |
| NC | 0.6 ± 0.1 | 1.3 ± 0.3 | 2.1 ± 0.4 | 0.8 ± 0.1 |
| CC | 5.0 ± 0.9<sup>a</sup> | 4.8 ± 0.9<sup>a</sup> | 7.4 ± 1.0<sup>a</sup> | 11.2 ± 2.9<sup>a</sup> |
| CT | 1.5 ± 0.1<sup>b</sup> | 1.4 ± 0.3<sup>b</sup> | 1.5 ± 0.2<sup>b</sup> | 1.7 ± 0.2<sup>b</sup> |
| NDV8 | 0.0 ± 0.0<sup>b</sup> | 0.0 ± 0.0<sup>b</sup> | 0.0 ± 0.0<sup>b</sup> | 0.0 ± 0.0<sup>b</sup> |
| NDV16 | 0.0 ± 0.0<sup>b</sup> | 0.0 ± 0.0<sup>b</sup> | 0.0 ± 0.0<sup>b</sup> | 0.0 ± 0.0<sup>b</sup> |
| NDV32 | 0.0 ± 0.0<sup>b</sup> | 0.0 ± 0.0<sup>b</sup> | 0.0 ± 0.0<sup>b</sup> | 0.0 ± 0.0<sup>b</sup> |
| NDV64 | 0.0 ± 0.0<sup>b</sup> | 0.0 ± 0.0<sup>b</sup> | 0.0 ± 0.0<sup>b</sup> | 0.0 ± 0.0<sup>b</sup> |
| CNDV8 | 0.0 ± 0.0<sup>b</sup> | 0.0 ± 0.0<sup>b</sup> | 1.0 ± 0.1<sup>b</sup> | 2.2 ± 0.0<sup>b</sup> |
| CNDV16 | 4.0 ± 0.2<sup>b</sup> | 2.3 ± 0.1<sup>b</sup> | 1.0 ± 0.1<sup>b</sup> | 3.3 ± 0.7<sup>b</sup> |
| CNDV32 | 3.7 ± 0.2<sup>b</sup> | 4.1 ± 0.2<sup>b</sup> | 3.3 ± 0.2<sup>b</sup> | 3.3 ± 0.2<sup>b</sup> |
| CNDV64 | 1.5 ± 0.2<sup>b</sup> | 0.5 ± 0.3<sup>b</sup> | 1.6 ± 0.3<sup>b</sup> | 2.3 ± 0.2<sup>b</sup> |
| CNDV8+T | 1.7 ± 0.1<sup>b</sup> | 0.8 ± 0.4<sup>b</sup> | 1.3 ± 0.4<sup>b</sup> | 0.0 ± 0.0<sup>b</sup> |
| CNDV16+T | 3.1 ± 0.1<sup>b</sup> | 1.8 ± 0.1<sup>b</sup> | 0.6 ± 0.1<sup>b</sup> | 0.0 ± 0.0<sup>b</sup> |
| CNDV32+T | 1.3 ± 0.1<sup>b</sup> | 1.2 ± 0.1<sup>b</sup> | 1.3 ± 0.1<sup>b</sup> | 0.0 ± 0.0<sup>b</sup> |
| CNDV64+T | 0.0 ± 0.0<sup>b</sup> | 0.6 ± 0.1<sup>b</sup> | 1.5 ± 0.1<sup>b</sup> | 3.0 ± 0.2<sup>b</sup> |
| NC | 1.0 ± 0.0 | 1.0 ± 0.0 | 1.0 ± 0.0 | 1.0 ± 0.0 |
| CC | 21.5 ± 12.0<sup>a</sup> | 29.5 ± 9.2<sup>a</sup> | 37.5 ± 7.8<sup>a</sup> | 56.5 ± 14.9<sup>a</sup> |
| CT | 24.5 ± 10.6 | 27.0 ± 8.5 | 1.0 ± 0.0<sup>b</sup> | 1.0 ± 0.0<sup>b</sup> |
| NDV8 | 1.0 ± 0.0<sup>b</sup> | 1.0 ± 0.0<sup>b</sup> | 1.0 ± 0.0<sup>b</sup> | 1.0 ± 0.0<sup>b</sup> |
| NDV16 | 1.0 ± 0.0<sup>b</sup> | 1.0 ± 0.0<sup>b</sup> | 7.5 ± 2.1<sup>b</sup> | 15.5 ± 3.5<sup>b</sup> |
| NDV32 | 1.0 ± 0.0<sup>b</sup> | 1.0 ± 0.0<sup>b</sup> | 1.0 ± 0.0<sup>b</sup> | 1.0 ± 0.0<sup>b</sup> |
| NDV64 | 1.0 ± 0.0<sup>b</sup> | 1.0 ± 0.0<sup>b</sup> | 1.0 ± 0.0<sup>b</sup> | 1.0 ± 0.0<sup>b</sup> |
| **CNDV8** | 12.5 ± 2.1<sup>b</sup> | 12.5 ± 3.5<sup>b</sup> | 13.0 ± 0.3<sup>b</sup> | 19.0 ± 1.4<sup>b</sup> |
| **CNDV16** | 21.5 ± 0.7 | 24.5 ± 3.5 | 15.5 ± 2.1<sup>b</sup> | 15.0 ± 1.4<sup>b</sup> |
| **CNDV32** | 15.0 ± 1.4 | 17.0 ± 0.0<sup>b</sup> | 15.5 ± 2.1<sup>b</sup> | 16.5 ± 2.1<sup>b</sup> |
| **CNDV64** | 16.0 ± 1.4 | 11.0 ± 4.2<sup>b</sup> | 12.5 ± 3.5<sup>b</sup> | 11.5 ± 3.5<sup>b</sup> |
| **CNDV8+T** | 11.0 ± 2.8<sup>b</sup> | 11.0 ± 4.2<sup>b</sup> | 14.5 ± 0.7<sup>b</sup> | 12.5 ± 3.5<sup>b</sup> |
| **CNDV16+T** | 13.0 ± 4.2<sup>b</sup> | 13.0 ± 5.6<sup>b</sup> | 13.5 ± 7.8<sup>b</sup> | 13.5 ± 2.1<sup>b</sup> |
| **CNDV32+T** | 11.5 ± 3.5<sup>b</sup> | 8.0 ± 2.8<sup>b</sup> | 12.0 ± 5.7<sup>b</sup> | 11.0 ± 5.7<sup>b</sup> |
| **CNDV64+T** | 9.5 ± 4.9<sup>b</sup> | 13.5 ± 4.9<sup>b</sup> | 18.5 ± 12.0<sup>b</sup> | 12.0 ± 4.2<sup>b</sup> |

**Table 6:**
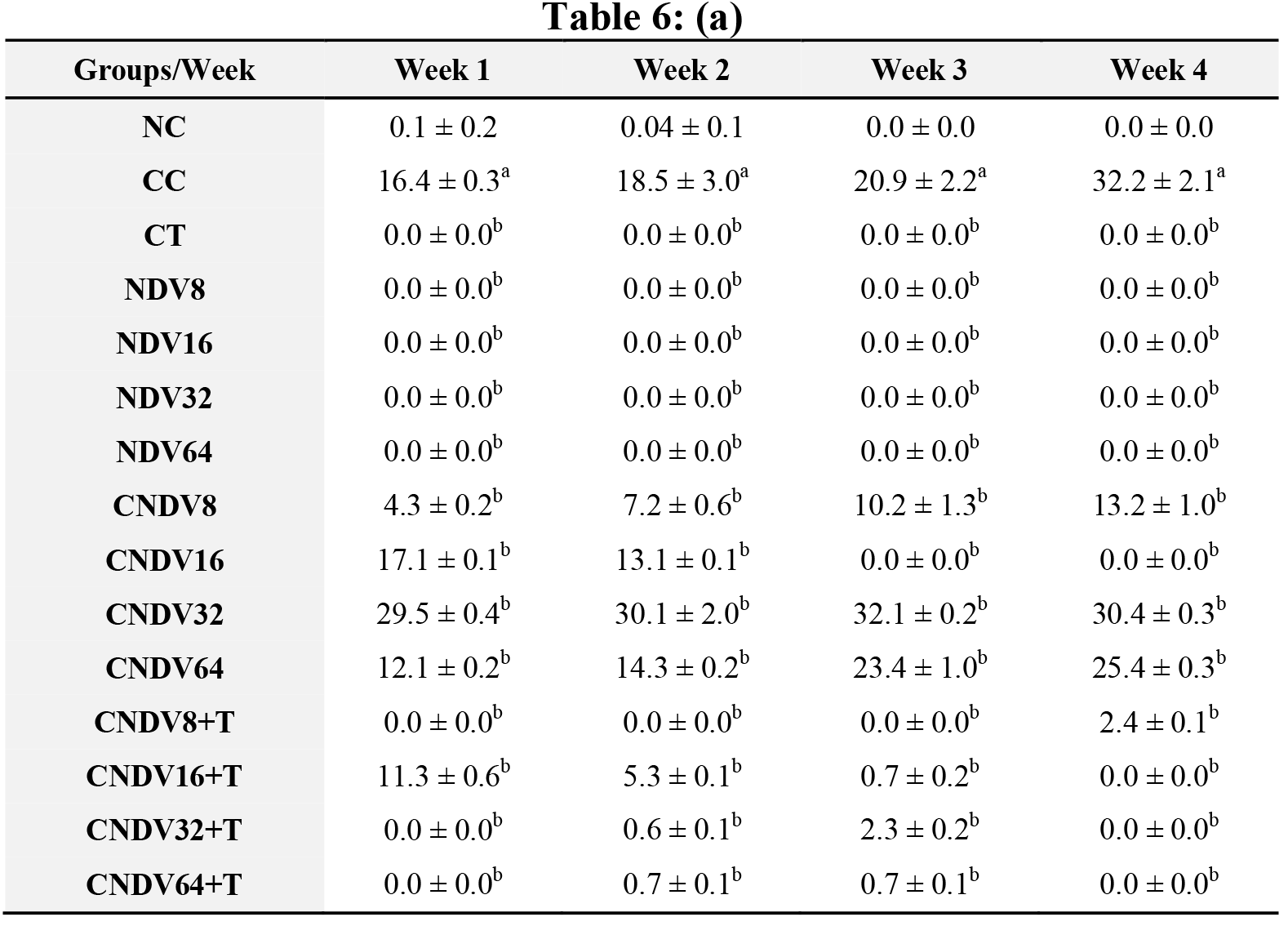
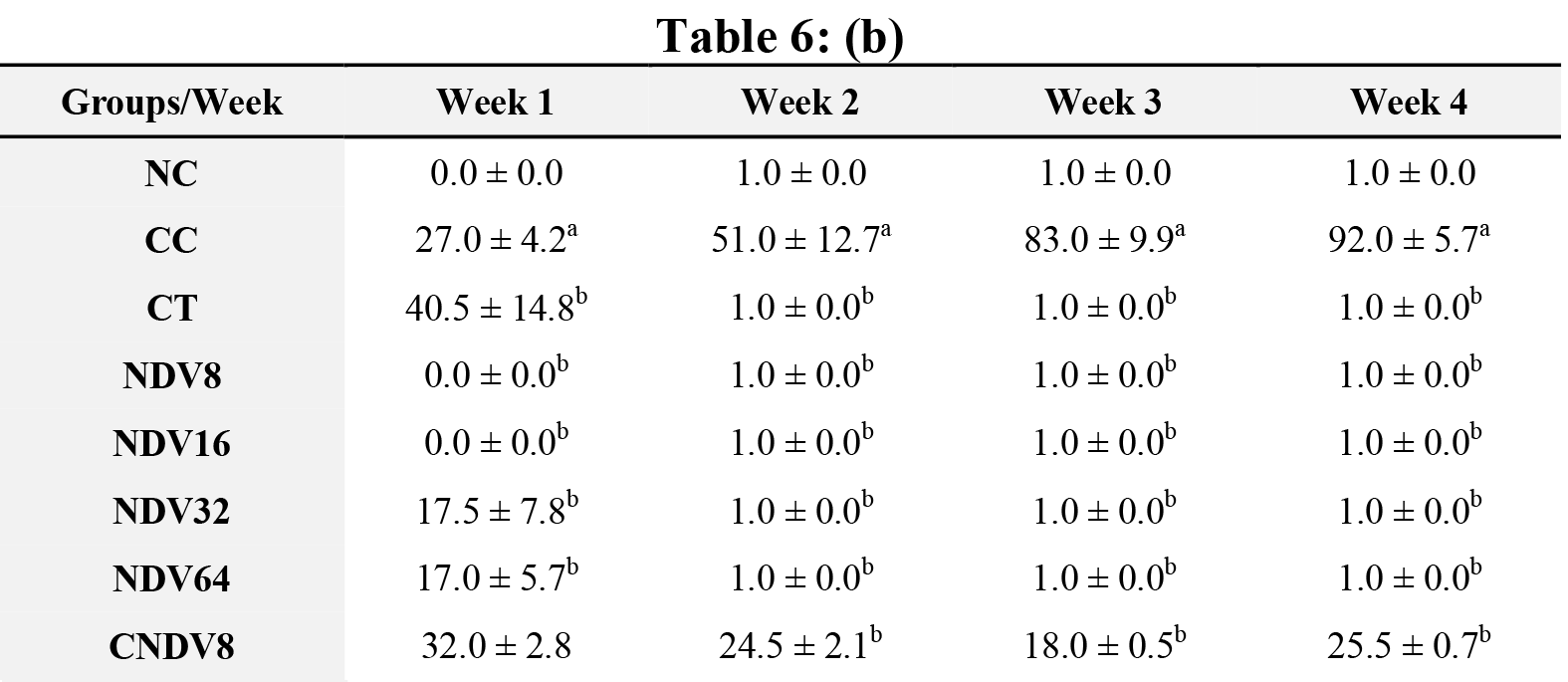
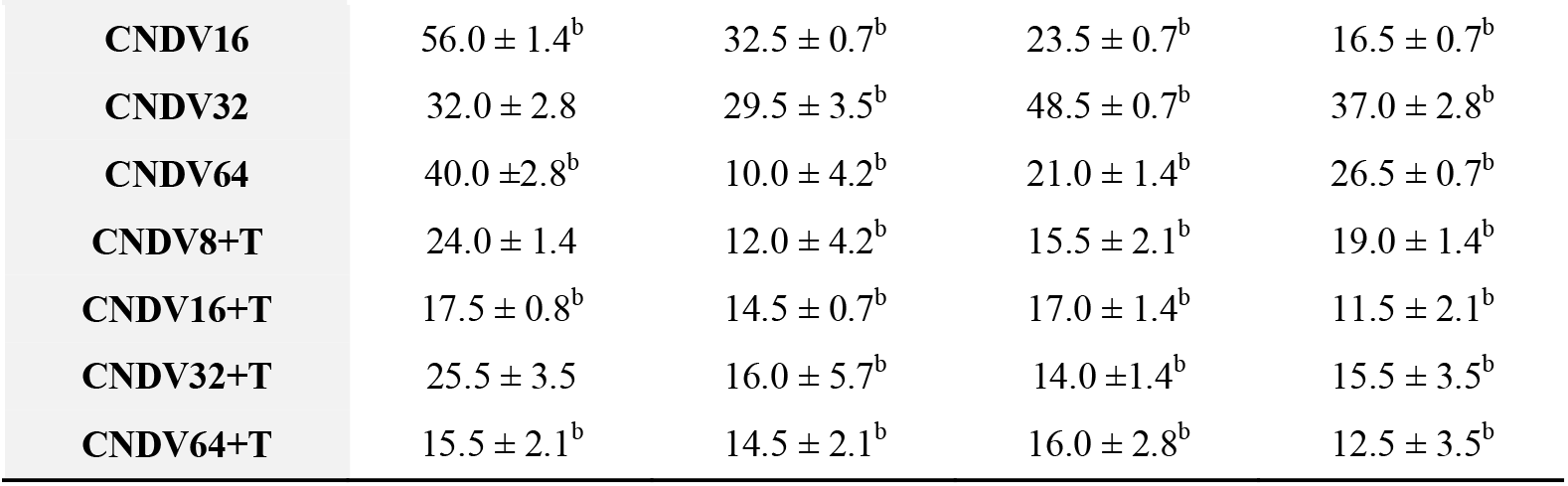
**(a)** Concentration (pg/ml) and **(b)** number spots of IL-10 in different groups of mice throughout four weeks experiment. NC: Normal Control; CC: Cancer Control; CT: Cancer + Tamoxifen; NDV: NDV virus alone; CNDV: Cancer and treated with NDV virus; CNDV + T: Cancer and treated with NDV and Tamoxifen. ^a^ *p* < 0.05 compared with NC; ^b^ *p* < 0.05 compared with CC.

| Groups/Week | Week 1 | Week 2 | Week 3 | Week 4 |
| --- | --- | --- | --- | --- |
| **NC** | 0.1 ± 0.2 | 0.04 ± 0.1 | 0.0 ± 0.0 | 0.0 ± 0.0 |
| **CC** | 16.4 ± 0.3<sup>a</sup> | 18.5 ± 3.0<sup>a</sup> | 20.9 ± 2.2<sup>a</sup> | 32.2 ± 2.1<sup>a</sup> |
| **CT** | 0.0 ± 0.0<sup>b</sup> | 0.0 ± 0.0<sup>b</sup> | 0.0 ± 0.0<sup>b</sup> | 0.0 ± 0.0<sup>b</sup> |
| **NDV8** | 0.0 ± 0.0<sup>b</sup> | 0.0 ± 0.0<sup>b</sup> | 0.0 ± 0.0<sup>b</sup> | 0.0 ± 0.0<sup>b</sup> |
| **NDV16** | 0.0 ± 0.0<sup>b</sup> | 0.0 ± 0.0<sup>b</sup> | 0.0 ± 0.0<sup>b</sup> | 0.0 ± 0.0<sup>b</sup> |
| **NDV32** | 0.0 ± 0.0<sup>b</sup> | 0.0 ± 0.0<sup>b</sup> | 0.0 ± 0.0<sup>b</sup> | 0.0 ± 0.0<sup>b</sup> |
| **NDV64** | 0.0 ± 0.0<sup>b</sup> | 0.0 ± 0.0<sup>b</sup> | 0.0 ± 0.0<sup>b</sup> | 0.0 ± 0.0<sup>b</sup> |
| **CNDV8** | 4.3 ± 0.2<sup>b</sup> | 7.2 ± 0.6<sup>b</sup> | 10.2 ± 1.3<sup>b</sup> | 13.2 ± 1.0<sup>b</sup> |
| **CNDV16** | 17.1 ± 0.1<sup>b</sup> | 13.1 ± 0.1<sup>b</sup> | 0.0 ± 0.0<sup>b</sup> | 0.0 ± 0.0<sup>b</sup> |
| **CNDV32** | 29.5 ± 0.4<sup>b</sup> | 30.1 ± 2.0<sup>b</sup> | 32.1 ± 0.2<sup>b</sup> | 30.4 ± 0.3<sup>b</sup> |
| **CNDV64** | 12.1 ± 0.2<sup>b</sup> | 14.3 ± 0.2<sup>b</sup> | 23.4 ± 1.0<sup>b</sup> | 25.4 ± 0.3<sup>b</sup> |
| **CNDV8+T** | 0.0 ± 0.0<sup>b</sup> | 0.0 ± 0.0<sup>b</sup> | 0.0 ± 0.0<sup>b</sup> | 2.4 ± 0.1<sup>b</sup> |
| **CNDV16+T** | 11.3 ± 0.6<sup>b</sup> | 5.3 ± 0.1<sup>b</sup> | 0.7 ± 0.2<sup>b</sup> | 0.0 ± 0.0<sup>b</sup> |
| **CNDV32+T** | 0.0 ± 0.0<sup>b</sup> | 0.6 ± 0.1<sup>b</sup> | 2.3 ± 0.2<sup>b</sup> | 0.0 ± 0.0<sup>b</sup> |
| **CNDV64+T** | 0.0 ± 0.0<sup>b</sup> | 0.7 ± 0.1<sup>b</sup> | 0.7 ± 0.1<sup>b</sup> | 0.0 ± 0.0<sup>b</sup> |
| **NC** | 0.0 ± 0.0 | 1.0 ± 0.0 | 1.0 ± 0.0 | 1.0 ± 0.0 |
| **CC** | 27.0 ± 4.2<sup>a</sup> | 51.0 ± 12.7<sup>a</sup> | 83.0 ± 9.9<sup>a</sup> | 92.0 ± 5.7<sup>a</sup> |
| **CT** | 40.5 ± 14.8<sup>b</sup> | 1.0 ± 0.0<sup>b</sup> | 1.0 ± 0.0<sup>b</sup> | 1.0 ± 0.0<sup>b</sup> |
| **NDV8** | 0.0 ± 0.0<sup>b</sup> | 1.0 ± 0.0<sup>b</sup> | 1.0 ± 0.0<sup>b</sup> | 1.0 ± 0.0<sup>b</sup> |
| **NDV16** | 0.0 ± 0.0<sup>b</sup> | 1.0 ± 0.0<sup>b</sup> | 1.0 ± 0.0<sup>b</sup> | 1.0 ± 0.0<sup>b</sup> |
| **NDV32** | 17.5 ± 7.8<sup>b</sup> | 1.0 ± 0.0<sup>b</sup> | 1.0 ± 0.0<sup>b</sup> | 1.0 ± 0.0<sup>b</sup> |
| **NDV64** | 17.0 ± 5.7<sup>b</sup> | 1.0 ± 0.0<sup>b</sup> | 1.0 ± 0.0<sup>b</sup> | 1.0 ± 0.0<sup>b</sup> |
| **CNDV8** | 32.0 ± 2.8 | 24.5 ± 2.1<sup>b</sup> | 18.0 ± 0.5<sup>b</sup> | 25.5 ± 0.7<sup>b</sup> |
| CNDV16 | 56.0 ± 1.4<sup>b</sup> | 32.5 ± 0.7<sup>b</sup> | 23.5 ± 0.7<sup>b</sup> | 16.5 ± 0.7<sup>b</sup> |
| CNDV32 | 32.0 ± 2.8 | 29.5 ± 3.5<sup>b</sup> | 48.5 ± 0.7<sup>b</sup> | 37.0 ± 2.8<sup>b</sup> |
| CNDV64 | 40.0 ± 2.8<sup>b</sup> | 10.0 ± 4.2<sup>b</sup> | 21.0 ± 1.4<sup>b</sup> | 26.5 ± 0.7<sup>b</sup> |
| CNDV8+T | 24.0 ± 1.4 | 12.0 ± 4.2<sup>b</sup> | 15.5 ± 2.1<sup>b</sup> | 19.0 ± 1.4<sup>b</sup> |
| CNDV16+T | 17.5 ± 0.8<sup>b</sup> | 14.5 ± 0.7<sup>b</sup> | 17.0 ± 1.4<sup>b</sup> | 11.5 ± 2.1<sup>b</sup> |
| CNDV32+T | 25.5 ± 3.5 | 16.0 ± 5.7<sup>b</sup> | 14.0 ± 1.4<sup>b</sup> | 15.5 ± 3.5<sup>b</sup> |
| CNDV64+T | 15.5 ± 2.1<sup>b</sup> | 14.5 ± 2.1<sup>b</sup> | 16.0 ± 2.8<sup>b</sup> | 12.5 ± 3.5<sup>b</sup> |

**Table 7:**
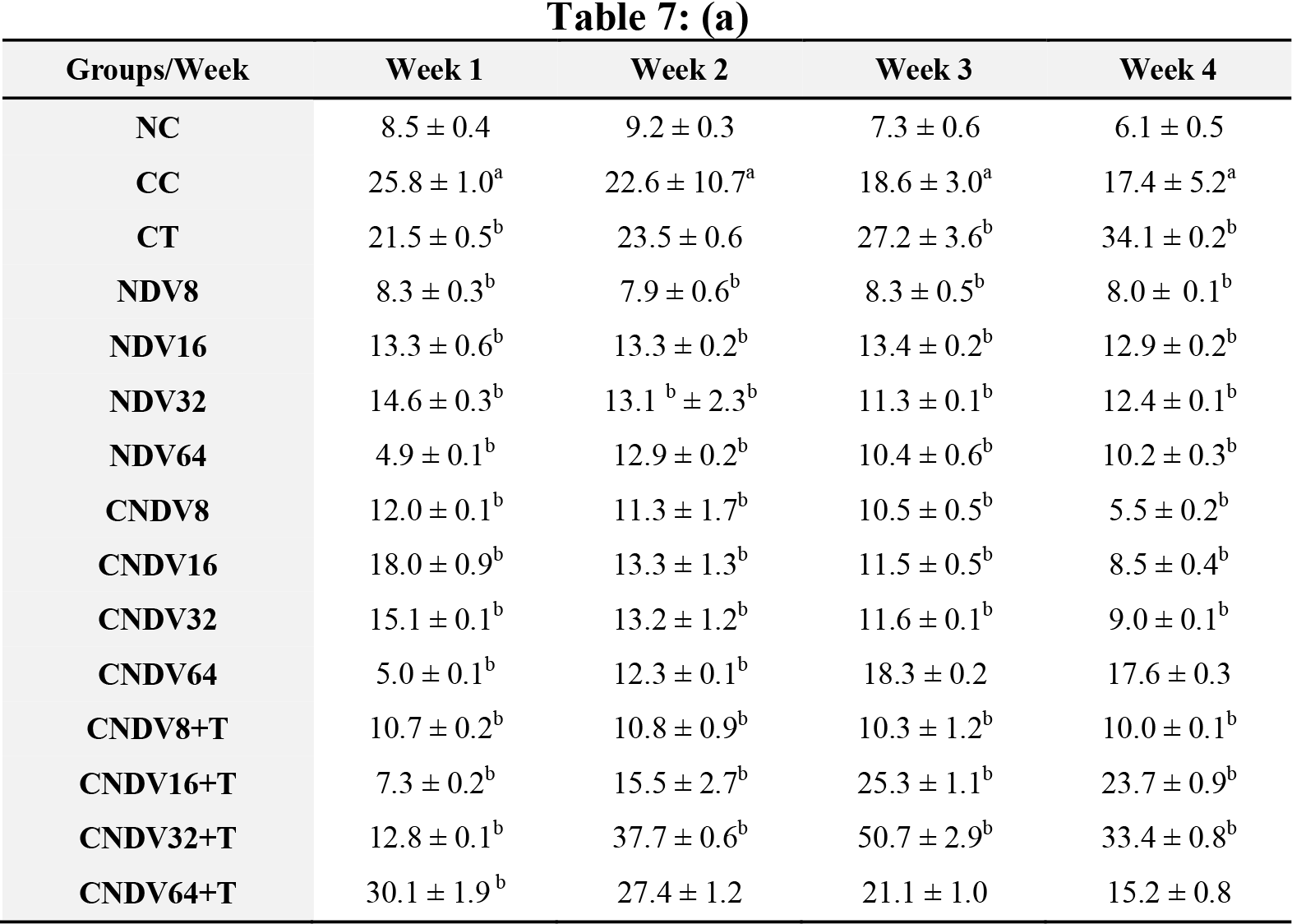
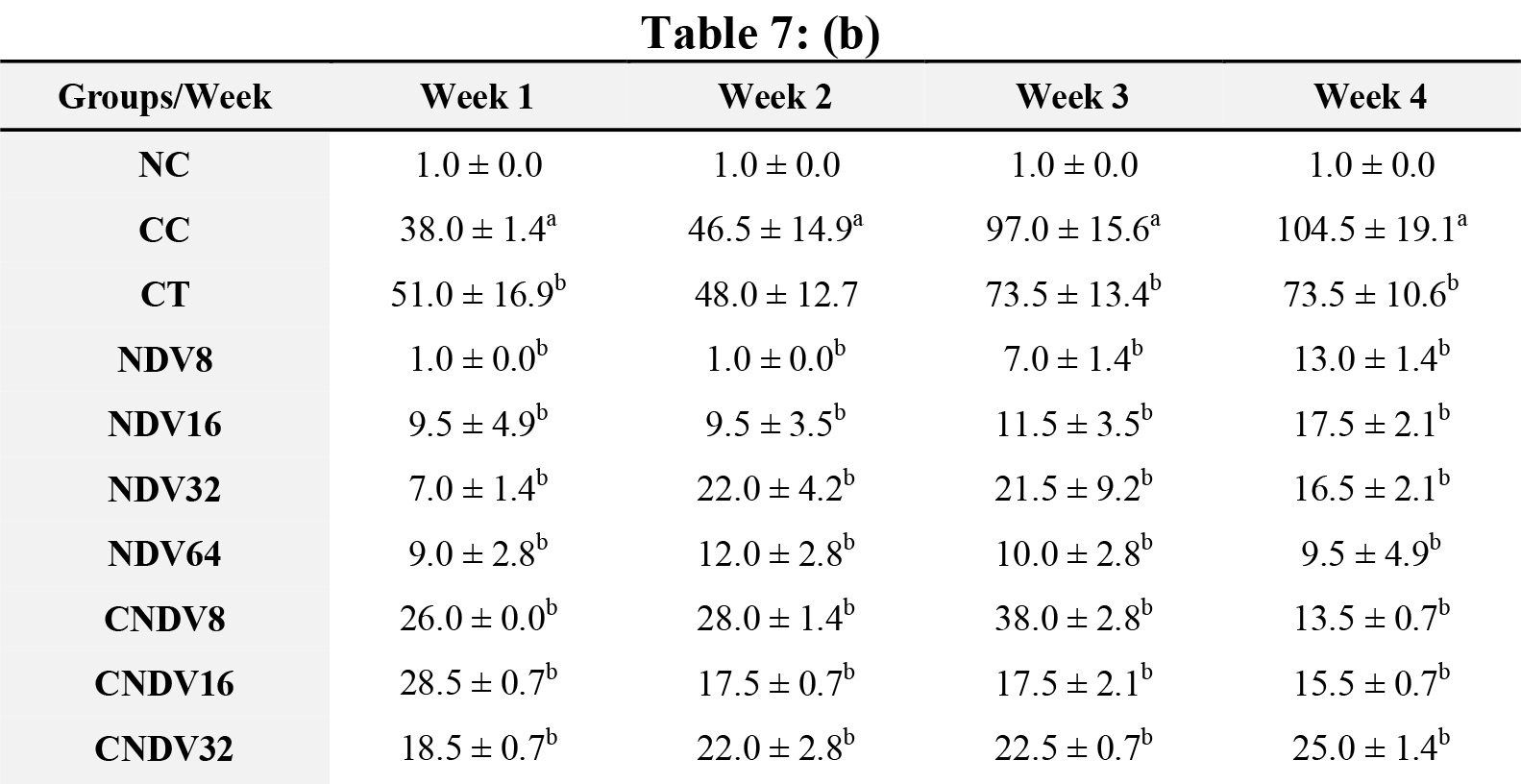
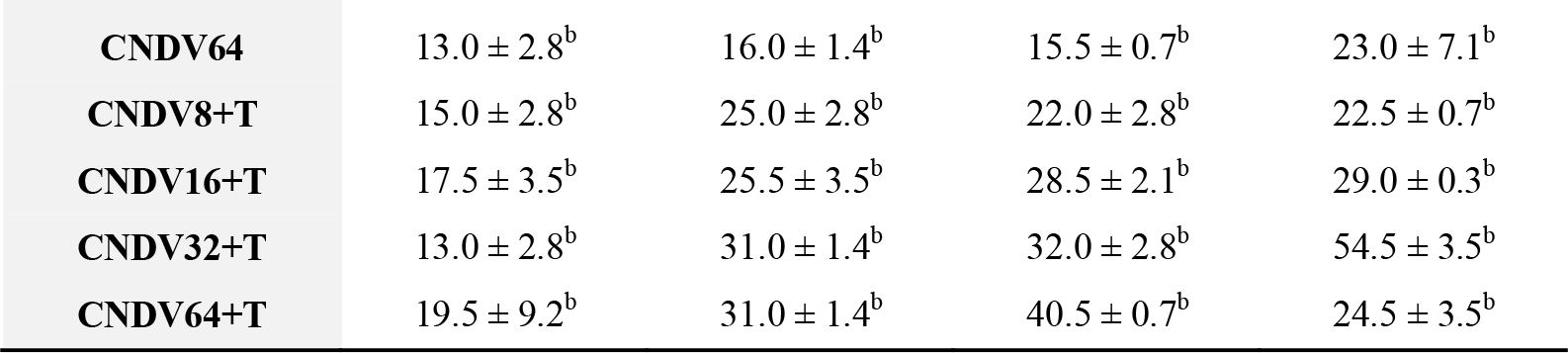
**(a)** Concentration (pg/ml) and **(b)** number spots of TNF-α in different groups of mice throughout four weeks experiment. NC: Normal Control; CC: Cancer Control; CT: Cancer + Tamoxifen; NDV: NDV virus alone; CNDV: Cancer and treated with NDV virus; CNDV + T: Cancer and treated with NDV and Tamoxifen. ^a^ *p* < 0.05 compared with NC; ^b^ *p* < 0.05 compared with CC.

| Groups/Week | Week 1 | Week 2 | Week 3 | Week 4 |
| --- | --- | --- | --- | --- |
| NC | 8.5 ± 0.4 | 9.2 ± 0.3 | 7.3 ± 0.6 | 6.1 ± 0.5 |
| CC | 25.8 ± 1.0<sup>a</sup> | 22.6 ± 10.7<sup>a</sup> | 18.6 ± 3.0<sup>a</sup> | 17.4 ± 5.2<sup>a</sup> |
| CT | 21.5 ± 0.5<sup>b</sup> | 23.5 ± 0.6 | 27.2 ± 3.6<sup>b</sup> | 34.1 ± 0.2<sup>b</sup> |
| NDV8 | 8.3 ± 0.3<sup>b</sup> | 7.9 ± 0.6<sup>b</sup> | 8.3 ± 0.5<sup>b</sup> | 8.0 ± 0.1<sup>b</sup> |
| NDV16 | 13.3 ± 0.6<sup>b</sup> | 13.3 ± 0.2<sup>b</sup> | 13.4 ± 0.2<sup>b</sup> | 12.9 ± 0.2<sup>b</sup> |
| NDV32 | 14.6 ± 0.3<sup>b</sup> | 13.1<sup>b</sup> ± 2.3<sup>b</sup> | 11.3 ± 0.1<sup>b</sup> | 12.4 ± 0.1<sup>b</sup> |
| NDV64 | 4.9 ± 0.1<sup>b</sup> | 12.9 ± 0.2<sup>b</sup> | 10.4 ± 0.6<sup>b</sup> | 10.2 ± 0.3<sup>b</sup> |
| CNDV8 | 12.0 ± 0.1<sup>b</sup> | 11.3 ± 1.7<sup>b</sup> | 10.5 ± 0.5<sup>b</sup> | 5.5 ± 0.2<sup>b</sup> |
| CNDV16 | 18.0 ± 0.9<sup>b</sup> | 13.3 ± 1.3<sup>b</sup> | 11.5 ± 0.5<sup>b</sup> | 8.5 ± 0.4<sup>b</sup> |
| CNDV32 | 15.1 ± 0.1<sup>b</sup> | 13.2 ± 1.2<sup>b</sup> | 11.6 ± 0.1<sup>b</sup> | 9.0 ± 0.1<sup>b</sup> |
| CNDV64 | 5.0 ± 0.1<sup>b</sup> | 12.3 ± 0.1<sup>b</sup> | 18.3 ± 0.2 | 17.6 ± 0.3 |
| CNDV8+T | 10.7 ± 0.2<sup>b</sup> | 10.8 ± 0.9<sup>b</sup> | 10.3 ± 1.2<sup>b</sup> | 10.0 ± 0.1<sup>b</sup> |
| CNDV16+T | 7.3 ± 0.2<sup>b</sup> | 15.5 ± 2.7<sup>b</sup> | 25.3 ± 1.1<sup>b</sup> | 23.7 ± 0.9<sup>b</sup> |
| CNDV32+T | 12.8 ± 0.1<sup>b</sup> | 37.7 ± 0.6<sup>b</sup> | 50.7 ± 2.9<sup>b</sup> | 33.4 ± 0.8<sup>b</sup> |
| CNDV64+T | 30.1 ± 1.9<sup>b</sup> | 27.4 ± 1.2 | 21.1 ± 1.0 | 15.2 ± 0.8 |
| NC | 1.0 ± 0.0 | 1.0 ± 0.0 | 1.0 ± 0.0 | 1.0 ± 0.0 |
| CC | 38.0 ± 1.4<sup>a</sup> | 46.5 ± 14.9<sup>a</sup> | 97.0 ± 15.6<sup>a</sup> | 104.5 ± 19.1<sup>a</sup> |
| CT | 51.0 ± 16.9<sup>b</sup> | 48.0 ± 12.7 | 73.5 ± 13.4<sup>b</sup> | 73.5 ± 10.6<sup>b</sup> |
| NDV8 | 1.0 ± 0.0<sup>b</sup> | 1.0 ± 0.0<sup>b</sup> | 7.0 ± 1.4<sup>b</sup> | 13.0 ± 1.4<sup>b</sup> |
| NDV16 | 9.5 ± 4.9<sup>b</sup> | 9.5 ± 3.5<sup>b</sup> | 11.5 ± 3.5<sup>b</sup> | 17.5 ± 2.1<sup>b</sup> |
| NDV32 | 7.0 ± 1.4<sup>b</sup> | 22.0 ± 4.2<sup>b</sup> | 21.5 ± 9.2<sup>b</sup> | 16.5 ± 2.1<sup>b</sup> |
| NDV64 | 9.0 ± 2.8<sup>b</sup> | 12.0 ± 2.8<sup>b</sup> | 10.0 ± 2.8<sup>b</sup> | 9.5 ± 4.9<sup>b</sup> |
| CNDV8 | 26.0 ± 0.0<sup>b</sup> | 28.0 ± 1.4<sup>b</sup> | 38.0 ± 2.8<sup>b</sup> | 13.5 ± 0.7<sup>b</sup> |
| CNDV16 | 28.5 ± 0.7<sup>b</sup> | 17.5 ± 0.7<sup>b</sup> | 17.5 ± 2.1<sup>b</sup> | 15.5 ± 0.7<sup>b</sup> |
| CNDV32 | 18.5 ± 0.7<sup>b</sup> | 22.0 ± 2.8<sup>b</sup> | 22.5 ± 0.7<sup>b</sup> | 25.0 ± 1.4<sup>b</sup> |
| **CNDV64** | 13.0 ± 2.8<sup>b</sup> | 16.0 ± 1.4<sup>b</sup> | 15.5 ± 0.7<sup>b</sup> | 23.0 ± 7.1<sup>b</sup> |
| **CNDV8+T** | 15.0 ± 2.8<sup>b</sup> | 25.0 ± 2.8<sup>b</sup> | 22.0 ± 2.8<sup>b</sup> | 22.5 ± 0.7<sup>b</sup> |
| **CNDV16+T** | 17.5 ± 3.5<sup>b</sup> | 25.5 ± 3.5<sup>b</sup> | 28.5 ± 2.1<sup>b</sup> | 29.0 ± 0.3<sup>b</sup> |
| **CNDV32+T** | 13.0 ± 2.8<sup>b</sup> | 31.0 ± 1.4<sup>b</sup> | 32.0 ± 2.8<sup>b</sup> | 54.5 ± 3.5<sup>b</sup> |
| **CNDV64+T** | 19.5 ± 9.2<sup>b</sup> | 31.0 ± 1.4<sup>b</sup> | 40.5 ± 0.7<sup>b</sup> | 24.5 ± 3.5<sup>b</sup> |

**Table 8:**
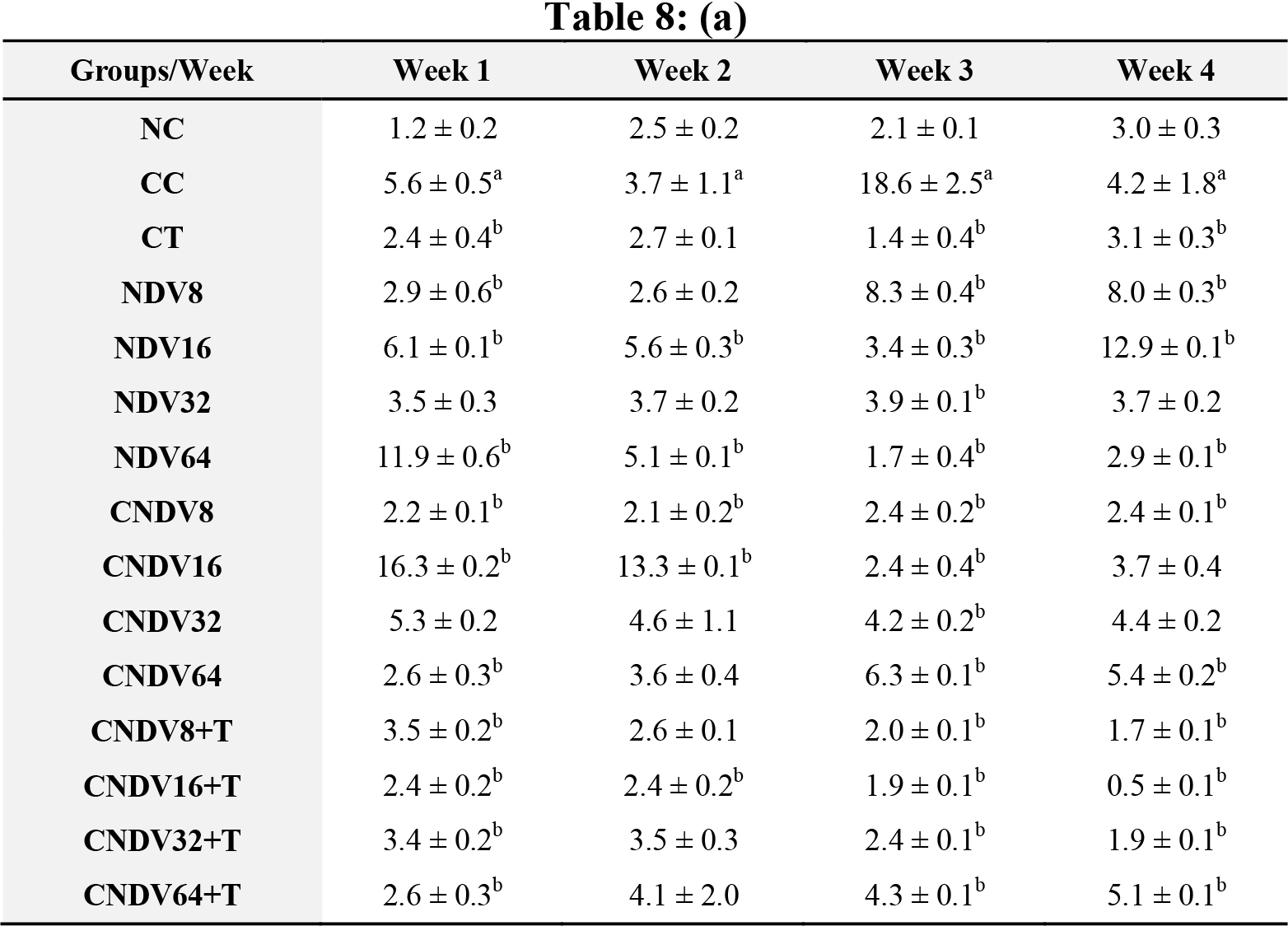
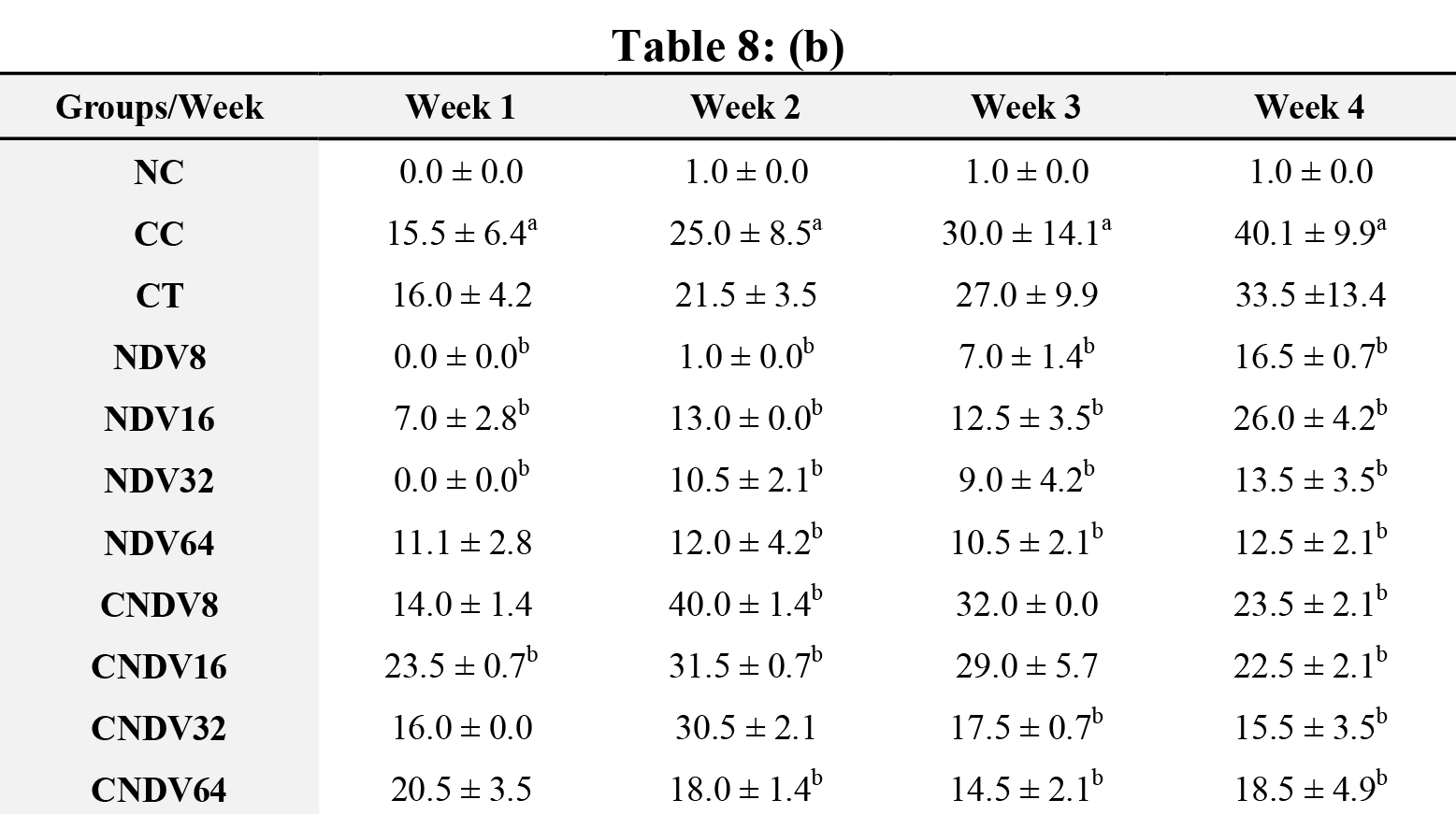
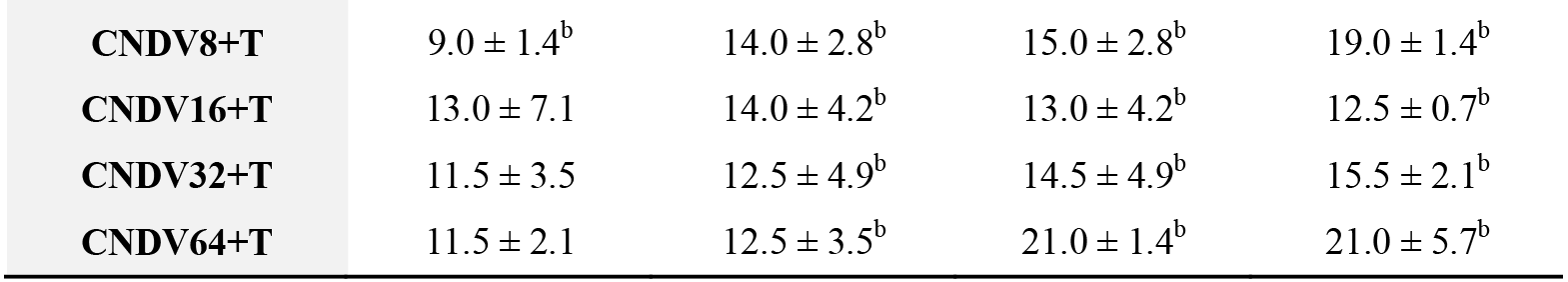
**(a)** Concentration (pg/ml) and **(b)** number spots of IFN-γ in different groups of mice throughout four weeks experiment. NC: Normal Control; CC: Cancer Control; CT: Cancer + Tamoxifen; NDV: NDV virus alone; CNDV: Cancer and treated with NDV virus; CNDV + T: Cancer and treated with NDV and Tamoxifen. ^a^ *p* < 0.05 compared with NC; ^b^ *p* < 0.05 compared with CC.

| Groups/Week | Week 1 | Week 2 | Week 3 | Week 4 |
| --- | --- | --- | --- | --- |
| NC | 1.2 ± 0.2 | 2.5 ± 0.2 | 2.1 ± 0.1 | 3.0 ± 0.3 |
| CC | 5.6 ± 0.5<sup>a</sup> | 3.7 ± 1.1<sup>a</sup> | 18.6 ± 2.5<sup>a</sup> | 4.2 ± 1.8<sup>a</sup> |
| CT | 2.4 ± 0.4<sup>b</sup> | 2.7 ± 0.1 | 1.4 ± 0.4<sup>b</sup> | 3.1 ± 0.3<sup>b</sup> |
| NDV8 | 2.9 ± 0.6<sup>b</sup> | 2.6 ± 0.2 | 8.3 ± 0.4<sup>b</sup> | 8.0 ± 0.3<sup>b</sup> |
| NDV16 | 6.1 ± 0.1<sup>b</sup> | 5.6 ± 0.3<sup>b</sup> | 3.4 ± 0.3<sup>b</sup> | 12.9 ± 0.1<sup>b</sup> |
| NDV32 | 3.5 ± 0.3 | 3.7 ± 0.2 | 3.9 ± 0.1<sup>b</sup> | 3.7 ± 0.2 |
| NDV64 | 11.9 ± 0.6<sup>b</sup> | 5.1 ± 0.1<sup>b</sup> | 1.7 ± 0.4<sup>b</sup> | 2.9 ± 0.1<sup>b</sup> |
| CNDV8 | 2.2 ± 0.1<sup>b</sup> | 2.1 ± 0.2<sup>b</sup> | 2.4 ± 0.2<sup>b</sup> | 2.4 ± 0.1<sup>b</sup> |
| CNDV16 | 16.3 ± 0.2<sup>b</sup> | 13.3 ± 0.1<sup>b</sup> | 2.4 ± 0.4<sup>b</sup> | 3.7 ± 0.4 |
| CNDV32 | 5.3 ± 0.2 | 4.6 ± 1.1 | 4.2 ± 0.2<sup>b</sup> | 4.4 ± 0.2 |
| CNDV64 | 2.6 ± 0.3<sup>b</sup> | 3.6 ± 0.4 | 6.3 ± 0.1<sup>b</sup> | 5.4 ± 0.2<sup>b</sup> |
| CNDV8+T | 3.5 ± 0.2<sup>b</sup> | 2.6 ± 0.1 | 2.0 ± 0.1<sup>b</sup> | 1.7 ± 0.1<sup>b</sup> |
| CNDV16+T | 2.4 ± 0.2<sup>b</sup> | 2.4 ± 0.2<sup>b</sup> | 1.9 ± 0.1<sup>b</sup> | 0.5 ± 0.1<sup>b</sup> |
| CNDV32+T | 3.4 ± 0.2<sup>b</sup> | 3.5 ± 0.3 | 2.4 ± 0.1<sup>b</sup> | 1.9 ± 0.1<sup>b</sup> |
| CNDV64+T | 2.6 ± 0.3<sup>b</sup> | 4.1 ± 2.0 | 4.3 ± 0.1<sup>b</sup> | 5.1 ± 0.1<sup>b</sup> |
| NC | 0.0 ± 0.0 | 1.0 ± 0.0 | 1.0 ± 0.0 | 1.0 ± 0.0 |
| CC | 15.5 ± 6.4<sup>a</sup> | 25.0 ± 8.5<sup>a</sup> | 30.0 ± 14.1<sup>a</sup> | 40.1 ± 9.9<sup>a</sup> |
| CT | 16.0 ± 4.2 | 21.5 ± 3.5 | 27.0 ± 9.9 | 33.5 ± 13.4 |
| NDV8 | 0.0 ± 0.0<sup>b</sup> | 1.0 ± 0.0<sup>b</sup> | 7.0 ± 1.4<sup>b</sup> | 16.5 ± 0.7<sup>b</sup> |
| NDV16 | 7.0 ± 2.8<sup>b</sup> | 13.0 ± 0.0<sup>b</sup> | 12.5 ± 3.5<sup>b</sup> | 26.0 ± 4.2<sup>b</sup> |
| NDV32 | 0.0 ± 0.0<sup>b</sup> | 10.5 ± 2.1<sup>b</sup> | 9.0 ± 4.2<sup>b</sup> | 13.5 ± 3.5<sup>b</sup> |
| NDV64 | 11.1 ± 2.8 | 12.0 ± 4.2<sup>b</sup> | 10.5 ± 2.1<sup>b</sup> | 12.5 ± 2.1<sup>b</sup> |
| CNDV8 | 14.0 ± 1.4 | 40.0 ± 1.4<sup>b</sup> | 32.0 ± 0.0 | 23.5 ± 2.1<sup>b</sup> |
| CNDV16 | 23.5 ± 0.7<sup>b</sup> | 31.5 ± 0.7<sup>b</sup> | 29.0 ± 5.7 | 22.5 ± 2.1<sup>b</sup> |
| CNDV32 | 16.0 ± 0.0 | 30.5 ± 2.1 | 17.5 ± 0.7<sup>b</sup> | 15.5 ± 3.5<sup>b</sup> |
| CNDV64 | 20.5 ± 3.5 | 18.0 ± 1.4<sup>b</sup> | 14.5 ± 2.1<sup>b</sup> | 18.5 ± 4.9<sup>b</sup> |
| CNDV8+T | 9.0 ± 1.4<sup>b</sup> | 14.0 ± 2.8<sup>b</sup> | 15.0 ± 2.8<sup>b</sup> | 19.0 ± 1.4<sup>b</sup> |
| CNDV16+T | 13.0 ± 7.1 | 14.0 ± 4.2<sup>b</sup> | 13.0 ± 4.2<sup>b</sup> | 12.5 ± 0.7<sup>b</sup> |
| CNDV32+T | 11.5 ± 3.5 | 12.5 ± 4.9<sup>b</sup> | 14.5 ± 4.9<sup>b</sup> | 15.5 ± 2.1<sup>b</sup> |
| CNDV64+T | 11.5 ± 2.1 | 12.5 ± 3.5<sup>b</sup> | 21.0 ± 1.4<sup>b</sup> | 21.0 ± 5.7<sup>b</sup> |

### 2.4. Detection of spleen produced-cytokine by ELISPOT

To determine the role of NDV virus induced – cytokines in suppressing 4T1 breast cancer, mice splenocytes cytokine production was measured after inoculation with NDV virus throughout the experimental period. Cancer control group (CC) possessed relatively high secretion of IL-6, IFN-γ, MCP-1, IL-10, IL-12p70 and TNF-α from week 1 until week 4 compared with other groups [Table 3(b) – 8(b)]. Groups that were treated with NDV virus alone did not show any significant difference compared with normal control (NC) group in all the cytokine measurement. In the case of all the transplanted category, group treated with virus alone (CNDV) secreted relatively high level of cytokines when compared with the group treated with combination of virus and tamoxifen (CNDV+T). On the other hand, there was no cytokines observed in normal control group.

## 3. Methods

### 3.1. Incubation of Embryonated Eggs

Propagation of NDV virus was performed based on the method reported by Blaskovis and Styk [27]. Embryonated chicken eggs aged 9 to 10 days were obtained from Linggi Poultry Farm, Negeri Sembilan, Malaysia. Upon arrival, the eggs were sprayed with 70% ethanol and wiped thoroughly with tissue paper to prevent contamination. The eggs were then kept in a 37°C-humidified incubator for 24 hours. The embryos were candled daily to monitor its viability. Same method was also used to determine the margin of air sac of the embryos, which were marked with a pencil prior to inoculation of the virus. All procedures were carried out under biological safety cabinet to minimise any contamination.

### 3.2. Seed Virus Dilution

Preparation of seed virus depends on the number of eggs used. Briefly, 10 fold of virus dilutions were prepared. First, three centrifuge tubes were filled with 9 ml of phosphate buffer saline (PBS) and the subsequent tubes were filled with 27 ml PBS. Approximately 1 ml of virus was filtered using 0.45 μM filter and added to the first centrifuge tube containing 9 ml PBS and suspended for several times to get 1 in 10 dilutions of the virus. By using a syringe, approximately 1 ml of the dilution was transferred to the second centrifuge tube before being suspended several times. This process was repeated until the third centrifuge tube. Finally, 3 ml of dilution from the third centrifuge tube was transferred to the fourth centrifuge tube containing 27 ml PBS to obtain a dilution of 10^-4^ NDV AF2240 which was used for the inoculation of virus in the embryonated chicken eggs.

### 3.3. Virus Inoculation

Virus was inoculated in eggs per method used by Alexander [28] with slight modification. A small hole approximately 1 mm in diameter was made using a sterile needle right above the air sac margin. By using a syringe, 0.1 ml of virus dilution was inoculated into each egg. Then the eggs were sterilised using 70% ethanol before using melted candle or sterile tape to seal the hole. The eggs were then kept inside the incubator and checked for dead embryos using candling after 48 hours. The eggs, which found to have dead embryos were removed and kept in the refrigerator at 4°C. The eggs were monitored daily for 96 hours or until 90% of the embryos died. All the eggs were kept in the refrigerator overnight to ensure that the blood vessel is constricted before virus harvesting process. This can avoid collection of blood during harvesting of the allantoic fluid.

### 3.4. Virus Harvesting

The eggs were left under a biological safety cabinet at room temperature for 30 minutes to avoid excessive condensation on the shells once removed from the refrigerator. The eggs shells above the air sac were then removed and the membranes were punctured to collect the allantoic fluid. If there were any visible contamination, the eggs were immediately rejected. To confirm the presence of NDV in the allantoic fluid, a rapid test using chicken red blood cells was conducted. The allantoic fluids collected were kept in sterile tubes. Immediately after all the allantoic fluids were harvested, the clarification and purification of the virus was carried out.

### 3.5. Virus Clarification and Purification

In brief, the clarification of allantoic fluid was done at 6000 g, 4°C for 10 minutes by using a refrigerated centrifuge. The supernatants were then centrifuged at 20,000 rpm, 4°C for 3 hours. Again, the supernatant was discarded while the pellet was re-suspended and dissolved in 1 ml NTE buffer (NaCl, Tri-HCl, EDTA). In addition, 30%, 40%, 50% and 60% of sucrose gradients were prepared in ultra-clear tubes and kept overnight at 4°C. A few drops of virus in NTE buffer were added to the sucrose solution by using a sterile pipette until all the tubes were equally balanced. The tubes were then centrifuged at 38,000 rpm, 4°C for 4 hours by using pre-cooled SW41 rotor. After the centrifugation, observation and marking of the purified band of virus was made under inverted microscope. The band was collected and transferred into polyalomer tubes. The tubes were topped-up with NTE buffer and balanced before subjected to centrifugation at 20,000 rpm at 4°C for 2 hours. The pellets obtained were dissolved in 1 ml NTE buffer and filtered using 0.4 μm filter. Finally, the suspensions were kept at −80°C until further use.

### 3.6. Preparation of Chicken Red Blood Cells for Virus Titration

Blood was withdrawn from the jugular vein of chicken by using syringe that filled with a mixture of PBS and EDTA to prevent the blood from clotting. The blood was transferred into 15 ml tube and topped-up with PBS, then centrifuged at 1000 rpm at room temperature for 10 minutes. The supernatant was discarded and the red blood cells were re-suspended in PBS and centrifuged again. This process was repeated for 3 more times. For virus titration purpose, 50 μl of the RBC was diluted in 100 ml PBS to get 0.5% suspension of RBC cell in PBS.

### 3.7. Haemagglutination (HA) Test

For HA test, 2^nd^ to 24^th^ well of 96-well plate were filled up with 50 μl of PBS while the 1^st^ well was filled up with 100 μl of purified virus. 50 μl of purified virus was transferred from the 1^st^ well into the 2^nd^ well to make a two-fold dilution and this continued until the 23^rd^ well. Then, 50 μl of the 0.5% RBC suspension was added into all wells and left for 30 minutes at room temperature. The 1^st^ well was served as positive control whereas the 24^th^ well served as negative control. Any appearance of red button was observed in all wells except the 24^th^ well that represented the virus HA titre.

### 3.8. NDV titre for Treatment

Viruses were prepared from titre of 10^8^ where it was neatly harvested from chicken eggs. Then the viruses were diluted to 8 HA, 16 HA, 32 HA and 64 HA unit of NDV as described in the Table 9 below: -

**Table 9:**
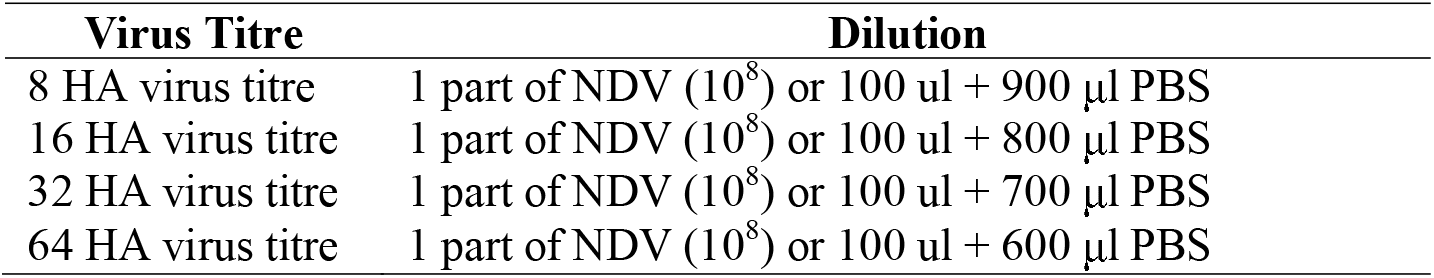
Showed preparation of virus titres from 10^8^ to get 8, 16, 32 and 64 HA units.

### 3.9. Cell Culture

Mouse mammary tumour cell line (4T1) was obtained from the American Type Culture Collection (ATCC). Cell was cultured in RPMI-1640 medium that was supplemented with 10% fetal bovine serum and 1% penicillin/streptomycin. Cells were maintained at 37°C in a humidified atmosphere of 5% CO_2_ in air. Culture medium was replaced every 2 to 3 days until the cell become 90% confluence before being subculture or used in the further experiment. For cancer cell induction purpose, the cell concentration was adjusted to 1 × 10^4^ cells/ml (0.1 cc injection per mice).

### 3.10. Animals

Healthy BALB/c mice weighing 15 – 20 g of age between 6-7 weeks old were obtained from Institute of Medical Research (IMR, Malaysia). The animals were reared in the animal house of Faculty of Medicine and Health Sciences, Universiti Putra Malaysia, where they were kept in plastic cages under hygienic conditions and were provided with standard animal feed. All the animal works were conducted in accordance with institutional guidelines for Animal Care and Use Committee (ACUC), Faculty of Medicine and Health Sciences, Universiti Putra Malaysia.

### 3.11. Experimental Design

Total 90 mice were used in this study and they divided into 15 groups (10 allotransplanted and 5 normal) with each group consisted of 6 mice. The normal category comprised of 5 groups of mice that were treated with 8, 16, 32 and 64 HA NDV without xenotransplant of cancer cells (NDV), while, a group that is only treated with normal feeding (without NDV and cancer cells) was served as control (NC). On the other hand, another 10 groups were allotransplanted with 4T1 breast cancer cells, which are under xenotransplantation category. Out of 10 groups, 4 groups were received 0.1 cc of 0.5 μg/ml Tamoxifen in combination with 8, 16, 32 and 64 HA of NDV, respectively (CNDV+T); another 4 groups were treated with 0.1 cc of 8, 16, 32 and 64 HA of NDV only (CNDV), respectively. A group of allotransplanted mice without subjected to any virus or tamoxifen was served as cancer control (CC), while, another group was only treated with 0.1 cc of 0.5 μg/ml tamoxifen was served as positive control (CT).

### 3.12. Tumour, Body Weight and Mortality

The body weight and tumour weight of the mice were measured on the first and last day after inoculation with NDV and allotransplanted with 4T1 breast cancer cells. The mortality rate of the mice for each group was also assessed every day until day 28 to determine the effect of NDV, NDV+Tamoxifen on the survival rate of mice.

### 3.13. Analysis of Liver Function

For assessment of liver function, plasma enzyme activities of total bilirubin level, alanine aminotransferase (ALT) and aspartate aminotransferase (AST) were determined using an automated procedure.

### 3.14. Cytokine Determination by Cytometric Bead Array (CBA)

Measurement of interleukin-6 (IL-6), interleukin-10 (IL-10), interleukin-12p70 (IL-12p70), interferon-γ (IFN-γ), monocyte chemoattractant protein-1 (MCP-1) and tumour necrosis factor-α (TNF-α) were performed using a mouse inflammation cytometric bead array kit (CBA; BD Biosciences, Malaysia). The assay protocol was done strictly according to the manufacturer’s instructions and the samples were analyzed with flow cytometer incorporated BD FACSComp^TM^ software.

### 3.15. Determination of Cytokines by Enzyme-Linked Immunosorbent Spot (ELISPOT)

Spleen cells suspension was prepared according to the method by Yang *et al*. [29] with slight modified. Spleen was removed and placed in plates containing Hank’s balanced salt solution (HBSS) under sterile techniques. Spleen was then cut into small pieces and mashed through a nylon sieve. Clumps and debris were removed by allowing them to settle down before the cell suspension was transferred to clean centrifuge tube. Cell suspension was then centrifuged at 200 g, 4°C for 15 minutes. Supernatant was discarded and cells were gently re-suspended with red cell lysing buffer in order to lyse erythrocytes. Again, the cell suspension was centrifuged at 200 g, 4°C for 5 minutes to remove cell debris and ghosts. After 2 washing in cold HBSS, the mononuclear cells were counted by trypan blue exclusion. Cell concentration was adjusted to the 1.5 to 3.0 × 10^5^ cells/ml in cell culture medium and kept on ice until further use. The ELISPOT assays were conducted by using the IL-6, IL-10, MCP-1, TNF-α, IFN-γ and IL-12p70 ELISPOT kit (BD Biosciences, Malaysia) according to the manufacturer’s protocol. Samples were run in quadruplicates. The numbers of spots were enumerated manually by inspection under a dissecting microscope and automated ELISPOT plate reader.

### 3.16. Statistical Analysis

Data are expressed as Mean±SD and analysis was done using Statistical Package for Social Sciences (SPSS version 21). Comparison between groups was determined by using ANOVA, and Dunnet test as *post-hoc* analysis. *p* < 0.05 was considered as significant.

## 4. Discussion

Virotherapy has emerged as a novel and potential anticancer agent in recent years. Numerous viruses were found to possess anti-cancer properties, especially oncolytic viruses[30–33]. Due to its unique mechanism of action and its ability to impact on many types of human cancer, NDV has been proposed as a potent cancer agent. Many strains of NDV were found to exhibit anti-cancer activity, for instance, MTH-68/H and AF2240 exhibited its anti-tumour effect via induction of nitric oxide synthesis from macrophages[34, 35]. In addition, systemic therapy with NDV strain PV701 showed a promising result as an important therapy for patients with a solid tumour [36]. Apart from that, other strains of NDV that have been investigated are HUJ [37], 73-T [38], V4UPM [39] and AF2240 [40].

There are two strains of NDV that can be found commonly in Malaysia, namely AF2240 and V4-UPM. In our previous studies, exposure of NDV AF2240 showed an apoptotic effect against breast tumour cells[23, 41]. However, the underlying mechanism of action on how NDV AF2240 kills the breast cancer cells remains largely unclear. In order to understand the mechanism of action, the current study aimed to evaluate the serological effects of NDV AF2240 inoculation in enhancing breast tumour by inhibiting cytokines secretion by the host immune system. In addition, the effect of NDV AF2240 on the liver was evaluated using liver function test throughout.

Based on *in vivo* pilot study conducted prior to this experiment, the minimum effective dosage required to suppress tumour cell by NDV AF2240 was 8HA (data not shown). Therefore, the dosage that was used in the current study started from 8HA up to 64HA. Due to this, the effect of high concentration of NDV AF2240 on body weight and mortality rate were investigated throughout the experiment. Interestingly, the results of the present study indicated that the NDV AF2240 was tolerated in mice even at 64HA. In addition, there were no significant adverse physical changes observed in the body weight and food intake of mice in the treated groups as compared to control group. Several studies reported that pre-and post-operative cancer patients show weight loss or cachexia [42, 43]. However, the overall body weight increased over time in the present study. Thus, it is suggested that the main contributor for weight increment was due to the tumour itself. In the case of tumour growth profile, tumour growth in all transplanted 4T1 breast cancer cells groups was completely inhibited by treatment with NDV alone. Conversely, low concentration of NDV+Tamoxifen (8 and 16HA) completely suppressed tumour growth, while at high concentrations (32 and 64HA), enhanced tumour cell growth was observed. Therefore, it is believed that there may have been some antagonistic effect between high titres of NDV with tamoxifen, neither allowing NDV nor tamoxifen to execute its effect in killing 4T1 tumour cells.

In order to evaluate the hepatotoxicity caused by the NDV as well as a combination of NDV and tamoxifen, liver biochemical parameters such as total bilirubin, AST and ALT were determined. Plasma concentrations of bilirubin, ALT and AST can be used as good indicators of the functionality and cellular integrity of the liver [44]. In addition, they are also good biomarkers that are used to predict any possible toxicity [45]. The present study demonstrated a significant elevation of bilirubin; AST and ALT in CNDV+T (32 and 64 HA) group compared to normal control. Moreover, the group treated with tamoxifen alone also showed significant increment in AST and ALT. This suggests that the high titre of NDV with tamoxifen or tamoxifen may cause damage to hepatocytes. Indeed, several studies revealed that that tamoxifen might cause toxic hepatitis, cirrhosis and sub massive hepatic necrosis [46, 47]. In terms of liver function enzymes, ALT is a more specific indicator of liver function, as compared to AST, which can also be found in red blood cells, cardiac and skeletal muscle [48]. Increased level of bilirubin may also indicate liver disease, biliary stricture, cardiac problem and neonatal hyperbilirubinaemia [49]. Therefore, careful consideration in interpreting the bilirubin level, ALT and AST enzyme activity has to be made with regards to findings from this study. Moreover, many factors can contribute to the elevation of the parameters observed here.

The current study showed that inoculation of NDV AF2240 could be beneficial for the treatment of breast cancer, but little is known about its mechanism of action. In order to elucidate a possible mechanism for the anti-neoplastic activity of NDV, immunological effects induced by NDV AF2240 was evaluated by determining the following cytokines; IL-6, IL-10, IL-12p70, IFN-γ, TNF-α and MCP-1. All the cytokines were measured by using ELISPOT and cytometric bead array assay. Many cytokines are known to be involved in inhibiting or enhancing tumour growth. For instance, IL-6, an inflammatory cytokine, is known to mediate many undesired, detrimental effects that contribute to cardiovascular disease [50], bowel disease [51] and Alzheimer disease [52]. In addition, IL-6 was found to pathologically regulate various types of cancer, including colorectal cancer [53], lung cancer [54] and breast cancer [55]. Recent studies reported that up-regulation of IL-6 level leads to cancer cells resistance to chemotherapeutic drugs [56]. It seems capable of protecting cells from damage by free radicals and this effect extends to the cells that might have escaped from normal cell cycle pathway [57]. Interestingly, findings reported here showed a significant suppression of IL-6 in the breast cancer-bearing mice challenged with NDV (8-64HA) and a combination of NDV and tamoxifen (8-64HA) compared with cancer control group (CC). Hence, NDV suppressed tumour growth may be through the impairment of IL-6 secretion.

As TNF-α is considered as a potential candidate for virotherapy in cancer patients, we have further analysed the expression level for TNF-α in NDV treated 4T1 breast cancer model. There are two mechanisms proposed for the antineoplastic activity of NDV: TNF-α secretion by activated peripheral blood mononuclear cells and enhancement in sensitivity of neoplastic cells towards TNF-α [58, 59]. In addition, recent studies also demonstrated that NDV induced apoptosis process mediated by TNF [60]. A part from that, it is also shown to enhance vascular permeability, facilitating the uptake and accumulation of chemotherapeutic drugs [61]. Hence, TNF-α therapy is a promising antineoplastic treatment. Present results revealed a marked increase of TNF-α concentration in 4T1 breast cancer bearing mice challenged with a combination of NDV Af2240 and tamoxifen. Although the result for cancer control is higher in CBA assay compared with ELISPOT, this may be that the cytokine-producing cell measurements done by using ELISPOT did not account for productivity per individual cells. Thus, it does not allow the differentiation on whether few cells produce many cytokines or many cells produce little cytokines as shown by CBA assay.

IL-12 is capable of evoking a potent immune response and also down-regulates the formation of new blood vessels into growing tumour [62]. IFN-γ is known as the key downstream factor induced by IL-12. Thus, the antitumour activity exerted by IL-12 is believed largely to be due to the local IFN-γ production and subsequent activation of the angiostatic chemokines IP-10 [63]. Nastala and her colleagues [64] reported that the tumour regression induced by recombinant IL-12 is associated with the production of IFN-γ. Hence, IL-12 therapy is crucial for cell-mediated-antitumour activity. However, results from this study revealed that treatment with NDV or NDV+Tamoxifen induces sustained low concentration of IL-12 and IFN-γ in breast cancer-bearing mice. This finding could suggest that NDV induces tumour regression via other signalling molecules or cytokines, but not IL-12 – IFN-γ pathway.

IL-10 is known to play an important role in inflammation. It suppresses the activation of macrophages and dendritic cells. In addition, it also inhibits numerous innate immune response and is associated with inflammatory activities [65]. On the other hand, high secretion of IL-10 has been found in various types of cancer cell, including breast, kidney, colon, lung and pancreas cancer [66–69]. In the context of breast cancer, IL-10 may act as a two-edged sword. Elevation of IL-10 level could facilitate the development of cancer by supporting tumour cell escape from host immune response via activation of PRL-R variants with altered pro-inflammatory or abrogated function. On the other hand, IL-10 could also prevent or reduce tumour growth and metastasis via suppression of angiogenesis [70]. Current results showed that cancer control (CC) group expressed a high level of IL-10, and this is similar to the studies done by Kozlowski *et al*. [71] and Razmkhah *et al*. [72]. Interestingly, breast cancer-bearing mice treated with a combination of NDV and tamoxifen showed a reduction in IL-10. Hence, it could be suggested that NDV+tamoxifen exhibited a synergistic effect, which prevents cancer growth through suppression of IL-10 secretion.

Lastly, the role of monocytes chemotactic protein-1 (MCP-1) was also evaluated in the current study. Several studies documented that upregulation of MCP-1 could assist in tumour growth by neovascularization [73, 74]. In addition, a clinical study done by Lebrecht *et al*. [75] reported that elevation of the MCP-1 level correlateds with advanced tumour stage and lymph node involvement in patients with breast cancer. Current data showed that there is only slight reduction of MCP-1 level in cancer bearing-mice treated with NDV alone or combination of NDV and tamoxifen compared to CC group. High concentration of MCP-1 found in CC could be due to release by cancer cells or inflammation [76]. Thus, NDV induced tumour regression may not be through suppression of MCP-1 pathway.

## 5. Conclusions

Based on the findings mentioned above, our data support that NDVAF2240 could potentially serve as an alternative treatment for breast cancer therapy. The dose or titre for virus needs proper research and study for its safety use and efficacy. A part from that, NDV AF2240 induced tumour regression may act through up-regulation or down-regulation of different cytokines. However, further fundamental research studies and data on tumour cytolysis *in vitro* and *in vivo* need to be done to discover the mechanism of action.

## Abbreviations

NDV: Newcastle disease virus
NC: Normal Control
CC: Cancer Control
CT: Cancer + Tamoxifen
CNDV: Cancer and treated with NDV virus
CNDV + T: Cancer and treated with NDV and Tamoxifen
IL-6: Interleukin 6
IFN-γ: Interferon gamma
MCP-1: Monocyte chemoattractant protein-1
IL-10: Interleukin 10
IL12p70: Interleukin 12
TNF-α: Tumour necrosis factor alpha

## 6. Declaration

### Ethics approval and consent to participate

This study was conducted in accordance with the recommendations from the guidelines provided by the Institutional Animal Care and Use Committee (IACUC) of the Faculty of Medicine and Health Sciences, Universiti Putra Malaysia. The protocol was approved and monitored by the Ethics Research Committee of the Universiti Putra Malaysia.

### Consent for publication

Not applicable

### Availability of data and material

Not applicable

### Competing of Interest

None of the authors have any competing interests.

### Funding

This work was supported by the Cancer Research Fund of Malaysia and GP-IPS University Research Grant provided to Prof. Dr. Fauziah Othman by National Cancer Council (MAKNA) and Universiti Putra Malaysia (UPM) respectively. Funding bodies did not have any influence in the design of the study and collection, analysis and interpretation of data or in writing the manuscript.

### Author’s Contributions

FO, JR, UA, ZE, AR and AI participated in the design of the research. JR carried out the experiments and analysed the data with the help from ZE and UA. YYK and UA analysed the data and wrote the paper. FO provided funding and supervised the study. All authors read and approved the final manuscript.

## Acknowledgments

We thank Dr. Mahmoud Bukar Maina of the University Sussex, UK for manuscript editing.

